# Expression of FoxP2 in the Basal Ganglia Regulates Vocal Motor Sequences in the Adult Songbird

**DOI:** 10.1101/2020.03.14.991042

**Authors:** Lei Xiao, Devin P. Merullo, Mou Cao, Marissa Co, Ashwinikumar Kulkarni, Genevieve Konopka, Todd F. Roberts

## Abstract

Disruption of the transcription factor FoxP2, which is enriched in the basal ganglia, impair vocal development in humans and songbirds. The basal ganglia are essential for the selection and sequencing of motor actions, but the circuit mechanisms governing accurate sequencing of learned vocalizations are unknown. Here, we show expression of FoxP2 in the basal ganglia is vital for the fluent initiation and termination of birdsong, and the maintenance of song syllable sequencing in adulthood. Knockdown of FoxP2 imbalances dopamine receptor expression across striatal direct-like and indirect-like pathways, suggesting a role of dopaminergic signaling in regulating vocal-motor sequencing. Confirming this prediction, we show that phasic dopamine activation, and not inhibition, during singing drives repetition of song syllables, thus also impairing fluent initiation and termination of birdsong. These findings demonstrate discrete circuit origins for the dysfluent repetition of vocal elements, a phenotype commonly observed in speech disorders.

## INTRODUCTION

Basal ganglia pathways are important for learning and controlling vocalizations^1, 2, 3^. The transcription factor forkhead box protein P2 (FOXP2) is enriched in medium spiny neurons (MSNs) in the striatal portion of the basal ganglia. Genetic disruptions of human *FOXP2* cause Childhood Apraxia of Speech, a developmental speech disorder marked by difficulties in controlling and accurately sequencing vocal-motor actions ^4, 5, 6^. FOXP2 has also been found to play a role in vocal learning in songbirds, indicating that it may have a common functional role in vocal-motor control across vocal learning species^2, 7, 8, 9, 10^. Nonetheless, how FOXP2 and basal ganglia circuits contribute to the fluent production of vocalizations is still poorly understood.

FoxP2 is strongly expressed in the songbird Area X, a specialized region of the striatum important for song learning and the control of song variability^11, 12, 13, 14, 15^. Its expression is thought to be necessary for learning song during development but not for the maintenance of song in adulthood. Knockdown of FoxP2 in Area X of juvenile zebra finches causes a variety of vocal deficits, including inaccurate syllable imitation, reduced stereotypy of song syntax, and anomalous repetition of song syllables^16^. In contrast to these effects on song development, FoxP2 appears to have a more limited, social context-dependent, role in adulthood. For instance, male zebra finches sing their song in a slightly more stereotyped manner when attempting to court a female bird^17, 18^. Knockdown of FoxP2 blocks this context-dependent decrease in song variability; however, it has not been reported to disrupt other aspects of syllable syntax, repetition of syllables, or the overall maintenance of learned song structure^19^.

FoxP2 is speculated to exert its effects in part by regulating dopamine signaling^7, 19, 20, 21^. Like the mammalian striatum, Area X receives glutamatergic input from pallial/cortical regions and input from dopamine neurons in the substantia nigra and ventral tegmental area (VTA). Disruptions of FoxP2 expression result in increased dopamine levels and reduced expression of dopamine receptors in the striatum^19, 20^. Consistent with developmental differences in the function of striatal FoxP2^16, 19^, dopamine signaling appears to also play a stronger role during song learning in juveniles than it does in adulthood. Ablating the dopaminergic input from VTA to Area X in juvenile birds results in inaccurate song imitation, whereas ablating dopaminergic input in adult birds has not been reported to disrupt the ability to accurately produce or maintain song ^22, 23^.

Nevertheless, several lines of evidence support the notion that basal ganglia circuits are still critical for adult song performance and song maintenance. Viral expression of the Huntington’s disease mutant gene fragment in Area X medium spiny neurons (MSNs) causes repetition of song syllables and disruptions in the sequencing of song ^24^. Neurotoxic lesions in Area X can also induce transient repetitions of song syllables^25, 26^. Lastly, deafening-induced deterioration of song is blocked by lesions of Area X, suggesting an active role for Area X in maintenance of adult song^13^.

Here, we apply novel reversible manipulations of gene expression and neural circuit activity to re-examine the role of striatal FoxP2 and dopaminergic circuits in the control of adult vocalizations. We find that FoxP2 is indeed critical for the fluent initiation and termination of adult song and regulation of song sequencing. Knockdown of FoxP2 using a small hairpin RNA (shRNA) reliably causes an increase in the repetition of song syllables. Turning off viral expression of the shRNA two months later rescues these disruptions in song. To clarify linkages between FoxP2 and dopaminergic circuitry, we mapped all Area X cell types and the distribution of FoxP2 and dopamine receptors using large-scale single-nucleus RNA sequencing. We find that FoxP2 knockdown causes a decrease in the expression of the D_1_ and D_5_ subtypes of dopamine receptors in Area X direct-like pathway MSNs. These results combined with previous studies indicating that mice carrying different substitutions in their endogenous *Foxp2* gene gave rise to specific alterations in dopamine levels in the striatum^7, 20^, encouraged us to test if manipulation of dopamine release in Area X altered song performance. We find that optogenetic excitation but not inhibition of VTA axon terminals in Area X during song production also disrupts the fluent initiation and termination of adult song, mimicking the most consistent effects of FoxP2 knockdown. Together, these findings indicate that FoxP2 regulated expression of dopamine receptors and changes in dopaminergic signaling interact in the striatum to control the fluent production of learned vocalizations.

## RESULTS

### Knockdown of FoxP2 Causes Progressive Vocal Repetitions and Disruptions in Syllable Sequencing

To drive a broad and genetically reversible knockdown of FoxP2 in adult zebra finch neurons, we employed a Cre-switch (CS) AAV platform^27^ (Figure 1a). A single transgene containing open reading frames of two fluorophores (mCherry and tagBFP), oriented in opposite directions, switches between expressing mCherry or BFP depending on Cre-driven recombination. Small hairpin RNAs against the zebra finch FoxP2 gene (shFoxP2) or scrambled hairpin (shScr) were inserted into the 3’UTR of the mCherry in the functional orientation with respect to the CAG promoter. In our constructs, shRNA is constitutively driven by the Pol II promoter CAG, and expression of the shRNA can therefore be turned off with introduction of Cre recombinase ^27, 28^.

**Figure 1.**
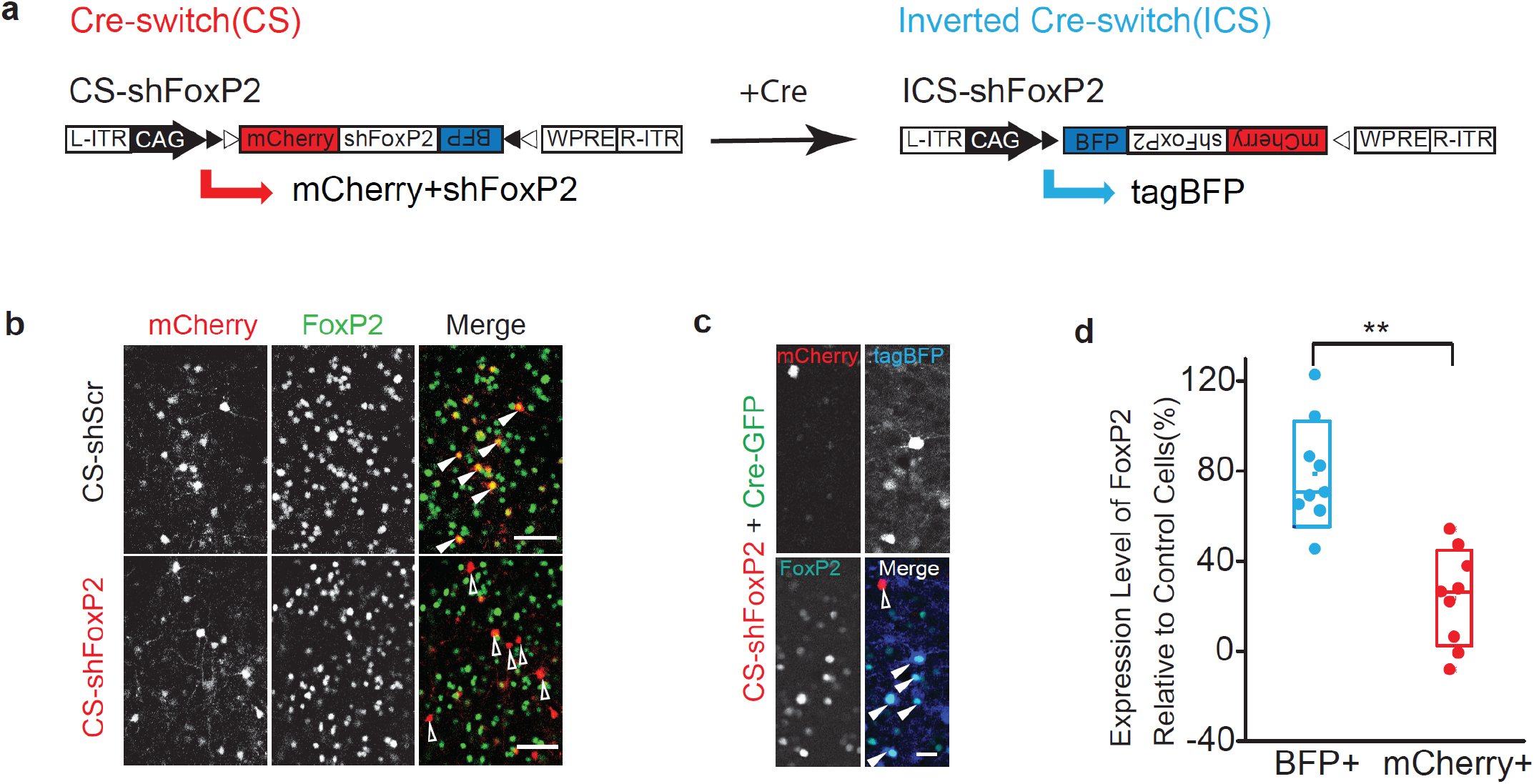
A Cre-switch Platform for Testing the Function of FoxP2 in Learned Vocalizations. (a) AAV constructs to achieve Cre-dependent silencing of zebra finch FoxP2. For this Cre-Switch (CS) configuration, the expression of mCherry and shRNAs are only maintained in the absence of Cre, whereas the expression of BFP is activated in the presence of Cre. WPRE, woodchuck polyresponse element. ITR, inverted terminal repeats. Open and filled triangles indicate loxP and lox2272, respectively. (b) FoxP2 expression in the cells infected with CS-shScr (top, filled triangles) and CS-shFoxP2 (bottom, open triangles) in Area X of adult birds. mCherry positive cells are cells infected with CS constructs. Scale bar, 50 μm. (c) confocal image shows that the expression of FoxP2 in mCherry+(open) and tagBFP+(filled) cells within Area X of an adult bird injected with CS-shFoxP2 and Cre-GFP constructs. Scale bar, 20 μm. (d) Quantification of the expression level of FoxP2 in mCherry+ and tagBFP+ cells (red 23.7±16.4% vs blue 78±18% relative to control cells) within Area X of adult birds injected with CS-shFoxP2 and Cre-GFP constructs (n=9 slices from 3 birds). The expression level of FoxP2 in tagBFP+ cells is significantly higher than in mCherry+ cells (p=0.0002, Mann-Whitney test). Box indicates the median ± 1.0 SD, mean shown by small dot.

We tested CS-shFoxP2 and CS-shScramble (CS-shScr) constructs *in vitro* and *in vivo* and found that we could significantly reduce the expression of FoxP2 following expression of CS-shFoxP2 (Figure 1b and S1 a-b). In contrast, the inverted Cre-switch (ICS) configuration of shFoxP2 resulted in no change in the expression of FoxP2 level *in vitro* (Figure S1c). Further testing of this configuration *in vitro* and *in vivo* found that the expression of FoxP2 was significantly higher in the neurons expressing ICS-shFoxP2 compared to those expressing CS-shFoxP2, suggesting that a second AAV expressing Cre-GFP could be used to rescue the knockdown of FoxP2 (Figures 1c, d and S1d). Upon the introduction of Cre, we observed a switch between mCherry and tagBGP fluorescence both *in vivo* and *in vitro* (Figure 1c and S1a,d-e), providing a reliable way to trace the knockdown and rescue efficiency as well as their coverage areas.

To test the effect of FoxP2 knockdown on vocal behavior, we made bilateral injections of CS-shFoxP2 into Area X of adult birds (159 ± 18 days post hatch (dph), n = 8 birds, Figure 2a) and monitored their song over two or more months (2.7±0.5 months). Adult zebra finch song is highly stereotyped and characterized by the precise ordering of individual song syllables^17^. We found that knockdown of FoxP2 in adult birds reliably disrupted this stereotypy in as little as three weeks following the viral injections. Overall, changes to song structure included anomalous repetition of individual syllables, replacement and deletion of some syllables from the song, and the creation and insertion of entirely new song syllables (Figures 2b-c, 3e and S2a). Some of these changes gave rise to significant alteration in the syntax of song and large-scale changes in how song bouts were produced.

**Figure 2.**
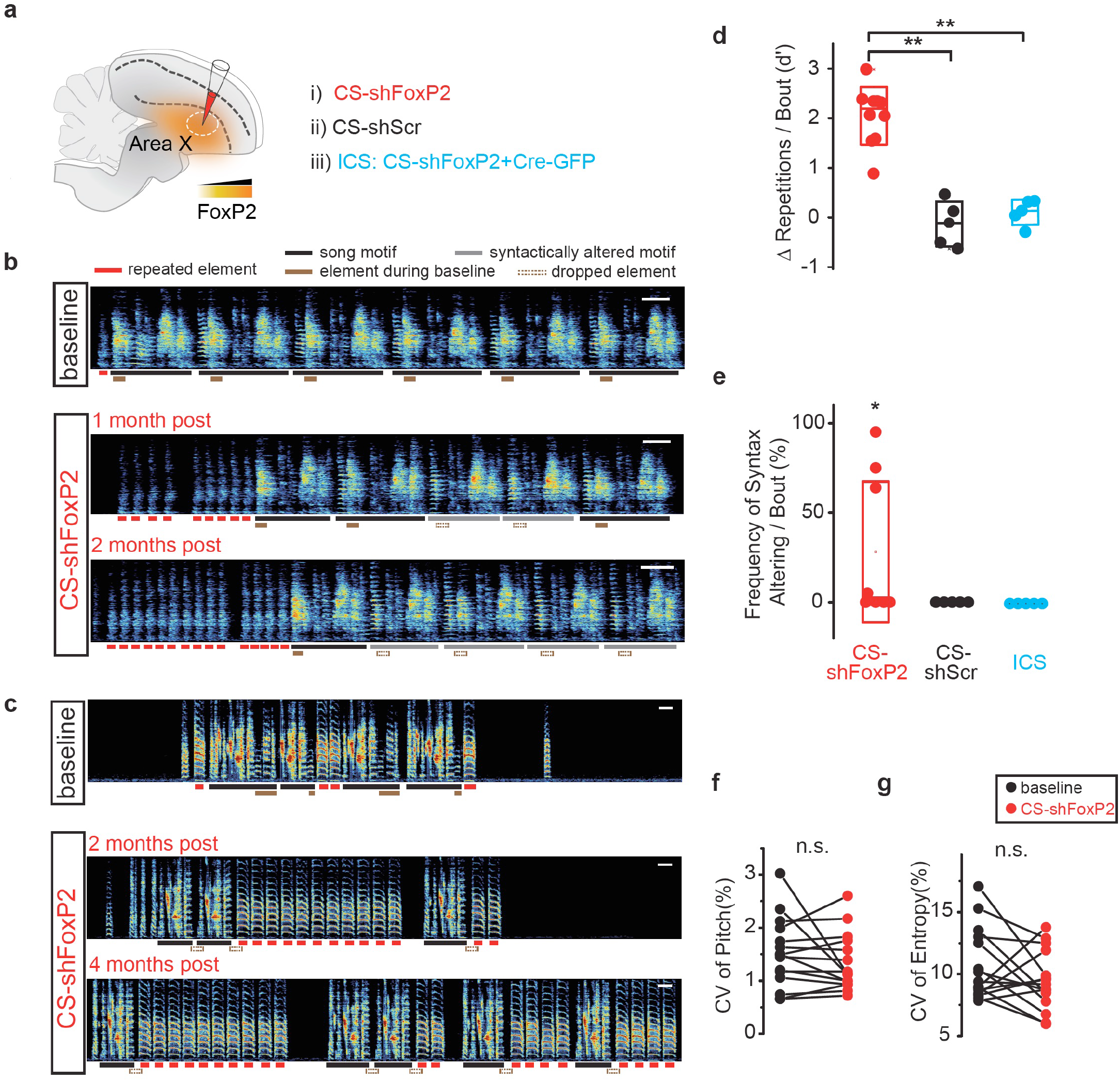
Diminished expression of FoxP2 in Area X Causes Vocal Repetitions and Disruptions in Syllable Sequencing in Adulthood. (a) CS constructs were bilaterally injected into Area X of adult birds, alone (i, CS-shFoxP2; ii, CS-shScr) or with Cre-GFP (iii, CS-shFoxP2/Cre-GFP, termed Inverted Cre Switch (ICS)). (b) Spectrograms of song recorded on the baseline day, 1 month, and 2 months after bilateral injection of CS-shFoxP2 construct in Area X of an adult bird. The number of repetitions of introductory elements in each song bout were increased over time, and one syllable was omitted over the course of 2 months. Scale bar, 200 ms. (c) Spectrograms of song recorded from a second bird on the baseline day, 2 months, and 4 months after bilaterally injection of CS-shFoxP2 construct in Area X. The number of repetitions of one syllable gradually increased over time, whereas other vocal elements were omitted over the course of 4 months. Scale bar, 200 ms. (d) Changes in the number of vocal repeats per song bout, expressed in units of *d*′, for CS-shFoxP2+ birds (red circles, *d*′=2.05±0.41, n=10 syllables from 8 birds), CS-shScr+ birds (black circles, *d*′= −0.13±0.56, n = 5 syllables from 5 birds), and ICS+ birds (blue circles, *d*′=0.11±0.31, n = 5 syllables from 5 birds). Changes in the number of repetitions of vocal elements in CS-ShFoxP2+ birds were significantly greater than changes observed in CS-shScr+ and ICS+ birds (CS-shScr, p= 0.0023; ICS, p=0.015, Kruskal-Wallis test). Box indicates the median ± 1.0 SD, mean shown by open dot. (e) Changes in the frequency of syntax altering per song bout for CS-shFoxP2+ birds (red circles, n=8 birds), CS-shScr+ birds (black circles, n = 5 birds), and ICS+ birds (blue circles, n = 5 birds). Change in the frequency of syntax altering in CS-ShFoxP2+ birds was significantly greater than zero (p=0.02, one tailed one sample t test). Box indicates the median ± 1.0 SD, mean shown by open dot. (f) Variability in pitch of syllables for baseline day (black, coefficient of variation [CV] = 1.52 ±0.17%) and two months post injection of CS-shFoxP2+ birds (red, CV = 1.34 ±0.14%). Knockdown of FoxP2 in Area X did not change the coefficient of variation of pitch (p = 0.75, n = 15, Wilcoxon matched-pairs signed-rank test). (g) Variability in entropy of syllables for baseline day (black, CV = 10.7 ±0. 75%) and two months post injection of CS-shFoxP2+ birds (red, CV = 9.46 ±0.62%). Knockdown of FoxP2 in Area X did not change the coefficient of variation of pitch (p = 0.17, n = 15, Wilcoxon matched-pairs signed-rank test).

Lesions of Area X can cause birds to transiently increase how many times they repeat introductory notes or other syllables that they already tended to repeat prior to lesions^25, 26^. We found significant increases in the repetition of previously repeated syllables at the beginning and/or end of song motifs in 7 out of 8 CS-shFoxP2+ birds (Figure 2b-d). FoxP2 knockdown birds appeared to get caught in ‘motor loops’, repeating a certain song syllable many times before transitioning to the next syllable. We also found that birds would begin to repeat syllables that were not repeated prior to FoxP2 knockdown (for example ‘green’ syllable in Figure 3e). In addition to increased repetition of introductory notes or pre-existing syllables at the end of the song motif, we found one bird also repeated syllable(s) that were created *de novo* following knockdown of FoxP2 (Figure S2a). In contrast to these changes in song, we did not detect changes in the number of syllable repetitions in any of the CS-shScr+ or ICS-shFoxP2+ birds (Figures 2a, d). These results suggest that FoxP2 in Area X helps regulate the initiation and termination of vocal sequences and can influence the transition from one syllable to the next within the song motif.

**Figure 3.**
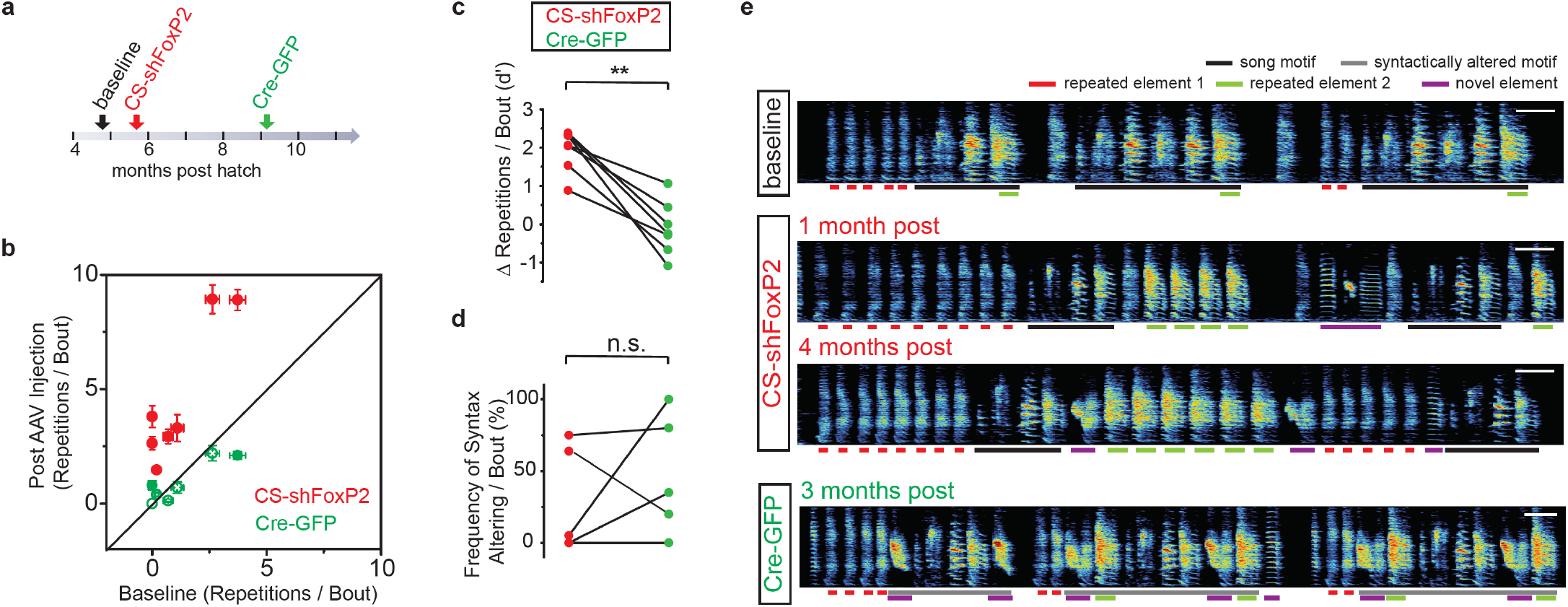
Reversal of FoxP2 Knockdown in Adult Zebra Finches Rescues Vocal Repetitions but not Disruptions in Song Syntax. (a) Strategy to rescue the expression levels of FoxP2 in Area X of adult birds. AAV encoding Cre-GFP was bilaterally injected into Area X 2-6 months after the initial knockdown of FoxP2 with CS-shFoxP2 construct. (b) Comparison of the number of repetitions of vocal element per song bout (± SEM) for the baseline day versus 2 months after injection of CS-shFoxP2 construct or 2 months after injection of Cre-GFP construct (n=7 vocal elements from 5 birds). Red filled circles represent vocal elements with significant differences between the baseline and post injection of CS-shFoxP2 (p<0.0001 and p=0.012, Kruskal-Wallis test). Green open circles represent vocal elements with nonsignificant differences (p>0.2, Kruskal- Wallis test), whereas green filled circles represent vocal elements with significant differences (p=0.014 and p=0.027, Kruskal-Wallis test) between the baseline and post injection of Cre-GFP. The number of repetitions of vocal elements after injection of Cre-GFP construct was significantly lower than the number of repetitions of the same elements after injection of CS-shFoxP2 construct (p = 0.016, n = 7, Wilcoxon matched-pairs signed-rank test). (c) Changes in the number of vocal repeats per song bout, expressed in units of *d*′, for CS-shFoxP2+ birds (n = 7 syllables 5 birds) between 2 months following CS-shFxoP2 injection (red circles, *d*′=1.94±0.21) and 3 months following Cre-GFP injection (green circles, *d*′=−0.011±0.27). Changes in the number of repetitions of vocal elements were significantly decreased following Cre-GFP injection (p=0.0007, paired t-test). (d) Changes in the frequency of syntax altering per song bout for CS-shFoxP2+ birds (n=5 birds) between 2 months following CS-shFxoP2 injection (red circles) and 3 months following Cre-GFP injection (green circles). Changes in the frequency of syntax altering were significantly greater than zero following Cre-GFP injection (p=0.033) but not following CS-shFoxP2 injection (p=0.084, one tailed one sample t-test). The frequencies of syntax altering were not significantly changed following Cre-GFP injection (p=0.38, paired t-test). (e) Spectrograms of song from one bird recorded on the baseline day, 1 month and 4 months after injection of CS-shFoxP2 construct, and 3 months after injection of Cre-GFP construct in Area X. Scale bar, 200 ms.

In addition to disrupting song linearity, we found syntax changes in 50% of CS-shFoxP2+ birds (4 of 8 birds, Figure 2e). These FoxP2 knockdown birds sang songs bouts with disrupted syntax a substantial proportion of the time (56.2 ± 19.4%), indicating that continued FoxP2 expression in Area X is important for the maintenance of learned syllable sequences and song structure. Most of the syntax changes arose from omitting one or multiple syllable(s) in the middle of the motif following FoxP2 knockdown (Figure 2b and S4a-b). In one bird we also observed both dropping of a pre-existing syllable and the addition of a new syllable to the bird’s song (Figure S2a and S3c). We did not detect these changes in any of the CS-shScr+ or ICS-shFoxP2+ birds which continued to sing their adult song normally after viral injections (0 of 10 birds; Figure 2e).

Disruptions to song structure observed in our FoxP2 knockdown birds could be the consequence of a more generalized disruption in song motor control, rather than a selective disruption in control of syllable transitions or syntax. Knockdown of FoxP2 increases the trial-by-trial variability of how individual syllables are produced and knockdown in adult birds disrupts social modulation of syllable variability^16, 19^. To examine if our FoxP2 knockdown approach disrupts trial-by-trial syllable variability, we quantified the coefficient of variation in syllable pitch and syllable entropy before and two months following FoxP2 knockdown. We did not find disruptions to syllable variability (Figures 2f-g), suggesting that the repetition and syntax changes observed in our knockdown experiments did not emerge from a general increase in motor variability. Together, these results demonstrate an essential and previously unappreciated role for FoxP2 in the selection and sequencing of previously learned vocal-motor actions.

### Rescue of FoxP2 Knockdown Reverses Disruptions in Syllable Repetitions but not Sequencing

Zebra finch song becomes less plastic and less reliant on sensory feedback during adulthood. For example, while deafening causes rapid deterioration of song in young adult birds, by ~180 days of age zebra finches can maintain a highly stable song for several weeks or months following deafening ^29, 30^. Given the large-scale changes to adult song observed following knockdown, we wondered if reversal of FoxP2 knockdown would result in recovery of the bird’s song.

We injected Area X with an AAV expressing Cre-GFP 2-6 months following injection of CS-shFoxP2 (AAVs expressing CS-shFoxP2 and Cre-GFP were injected at 177±24 dph and 277±23 dph, respectively, Figure 3a). We found that reversal of FoxP2 knockdown largely eliminated the increased repetition of introductory notes and song syllables within 3 months following injection of Cre-GFP (Figures 3b-c, e and S2a-b). In contrast, the changes in the syntax were either largely preserved or became more severe in three birds following injection of Cre-GFP (Figure 3d and S3a-e). While the dropping of syllables was prominent following injection of CS-shFoxP2, we found that the incorporation of new vocal elements was prominent during the song recovery period following injection of Cre-GFP (Figures S3a-e and S4a-e). Of the two CS-shFoxP2+ birds who did not show any syntax changes following injection of CS-shFoxP2, we found that in one, the frequency of syllable insertions per bout increased by 35% following reversal of the knockdown, while the other bird did not show any change in syntax throughout the time course of our experiment (Figures S3b, d). Only one of the five birds undergoing reversal of FoxP2 knockdown showed a decrease in syllable omitting following injection of Cre-GFP (Figures S2b and 3d). Together, these results indicate that reversal of FoxP2 knockdown is sufficient to return syllable repetition rates back to those observed prior to FoxP2 knockdown, but also resulted in birds ultimately singing songs that could be different from their pre-knockdown song due to the sustained omission of pre-existing syllables and/or the addition of a new syllable(s) into their song.

### Single-Nuclei RNA Sequencing Reveals Effects of FoxP2 Knockdown

FoxP2 is broadly expressed in Area X^15^ and, as a transcription factor, can have broad effects on gene expression. To identify subpopulation(s) of cells most affected by FoxP2 knockdown and the potential cellular mechanisms for disruptions in song fluency, we performed single-nuclei RNA (snRNA) sequencing on tissue samples of Area X from CS-shScr+ birds and CS-shFoxP2+ birds (for each group, 4 hemispheres total from 2 individuals). A combined clustering analysis indicated that FoxP2 knockdown did not significantly alter the distribution or composition of cell types in Area X (Figure 4a, Figure S5a), and cells from each group were well-represented within each cluster (Figure 4b).

**Figure 4.**
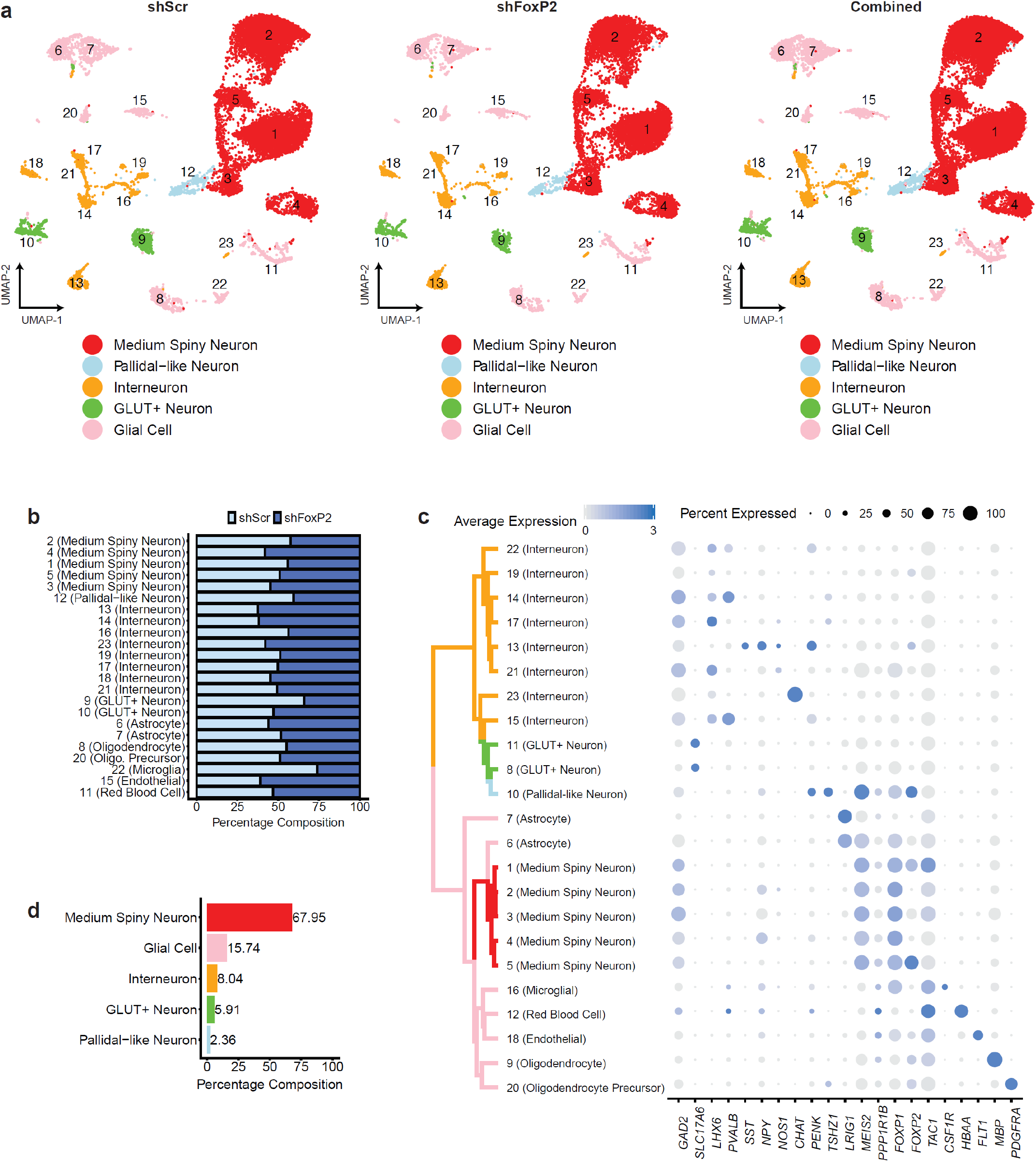
Overview of Cell Types in Area X (next page). (a) UMAP projections of nuclei from Area X (left, shScr subset; middle, shFoxP2 subset; right, combined analysis). Clusters are numbered in ascending order by decreasing size (1-largest; 23-smallest). (b) Cluster composition by each dataset (shScr or shFoxP2). (c) Hierarchical clustering and cell-type gene marker expression of an independent analysis of the shScr group. For marker gene expression, the size of the dot indicates the percent of nuclei within a cluster expressing a given gene, and the color of the dot indicates the average normalized expression level. (d) Overall cell type composition in Area X of the shScr group.

As a first catalogue of the cell type identities in Area X, we focused on an independent clustering analysis of the CS-shScr+ birds (Figure S5b) that identified 23 distinct cell groups (Figure 4c). Most cells were MSNs (identified by the strong collective expression of *Gad2*, *Ppp1r1b* ^31, 32^, and *FoxP1* ^33^, with five MSN clusters comprising 68% of total cells in Area X (Figure 4d). The pallidal-like cells, identified primarily by the expression of *Penk* ^31^, existed within one cluster and were far less numerous at only 2.4% of all cells (Figure 4d). Gene markers for the GPi ^32, 34^ and GPe ^35, 36^ were largely absent from any cluster in the dataset (Figure S6), although the pallidal-like cells here contain some similarities to arkypallidal cells reported in mammals (unpublished observations). We observed eight distinct clusters of GABAergic neurons, some of which likely correspond to known classes of striatal interneurons such as *Pvalb*+ interneurons, *Sst*+/*Npy*+/*Nos1*+ interneurons, and *Chat*+ interneurons ^31, 37, 38, 39, 40^. Two clusters expressed the glutamatergic transporter gene *Slc17a6*, which could relate to a glutamatergic cell-type that has been reported to exist in Area X ^41^. The remaining clusters comprised various glial cell-types such as astrocytes and oligodendrocytes (Figure 4c).

MSNs in Area X corresponding to the mammalian direct and indirect pathways MSNs have not been previously identified ^42^. Canonically, direct pathway MSNs are *Drd1*+ and *FoxP2*+, and project to the GPi while indirect pathway MSNs are *Drd2*+ and *FoxP2*−, and project to the GPe ^43, 44, 45, 46^. Area X MSNs, on the other hand, have been thought to be a more homogenous population expressing both *Drd1* and *Drd2,* and *FoxP2* ^47, 48^. Our snRNA-seq analysis of the CS-shScr+ birds revealed that 36% of the 9,672 MSNs are exclusively *Drd1*+ and/or *Drd5*+ (notated here as *Drd1/5+)*, while 13% are exclusively *Drd2*+. An additional 18% express varying levels of *Drd1/5* and *Drd2* consistent with previous studies. All five MSN clusters identified in our snRNA-seq dataset contain cells expressing *Drd1/5*, while only three contain cells expressing *Drd2* (Figure 5b). On the basis of this separation of *Drd2*+ cells across clusters, we performed a differential gene expression analysis by grouping the two clusters containing *Drd1*+/*Drd2*− cells into a putative direct-like pathway and the three clusters containing *Drd1*+ cells and *Drd2*+ cells (separate cells, not necessarily co-localizing *Drd1* and *Drd2*) into a putative indirect-like pathway. The direct-like pathway grouping significantly expressed genes that canonically mark direct pathway MSNs, including *FoxP2*, whereas the indirect-like pathway grouping significantly expressed genes that mark indirect pathway MSNs ^44, 46, 49^ (Figure 5c). Similar to what has been described in the mammalian direct and indirect pathways, *FoxP2* frequently co-localized with *Drd1*, but not with *Drd2* expressing neurons (Figures 5d-e). Of all cells that were *Drd1/5*+ and *Drd2*−, 61% also expressed *FoxP2*. However, of cells that were *Drd1/5*− and *Drd2*+, only 21% co-expressed *FoxP2* (Figure 5f). Thus, MSNs in Area X can segregate into broad classes corresponding to direct-like (*Drd1*+/*FoxP2*+) and indirect-like pathways (*Drd2*+/*FoxP2*−) (Figure 5a), as well as a third *Drd1*/2+ class. Further, these classes map to different clusters based on more restrictive expression patterns of *Drd2* and *FoxP2* (Figures 5b-f).

**Figure 5.**
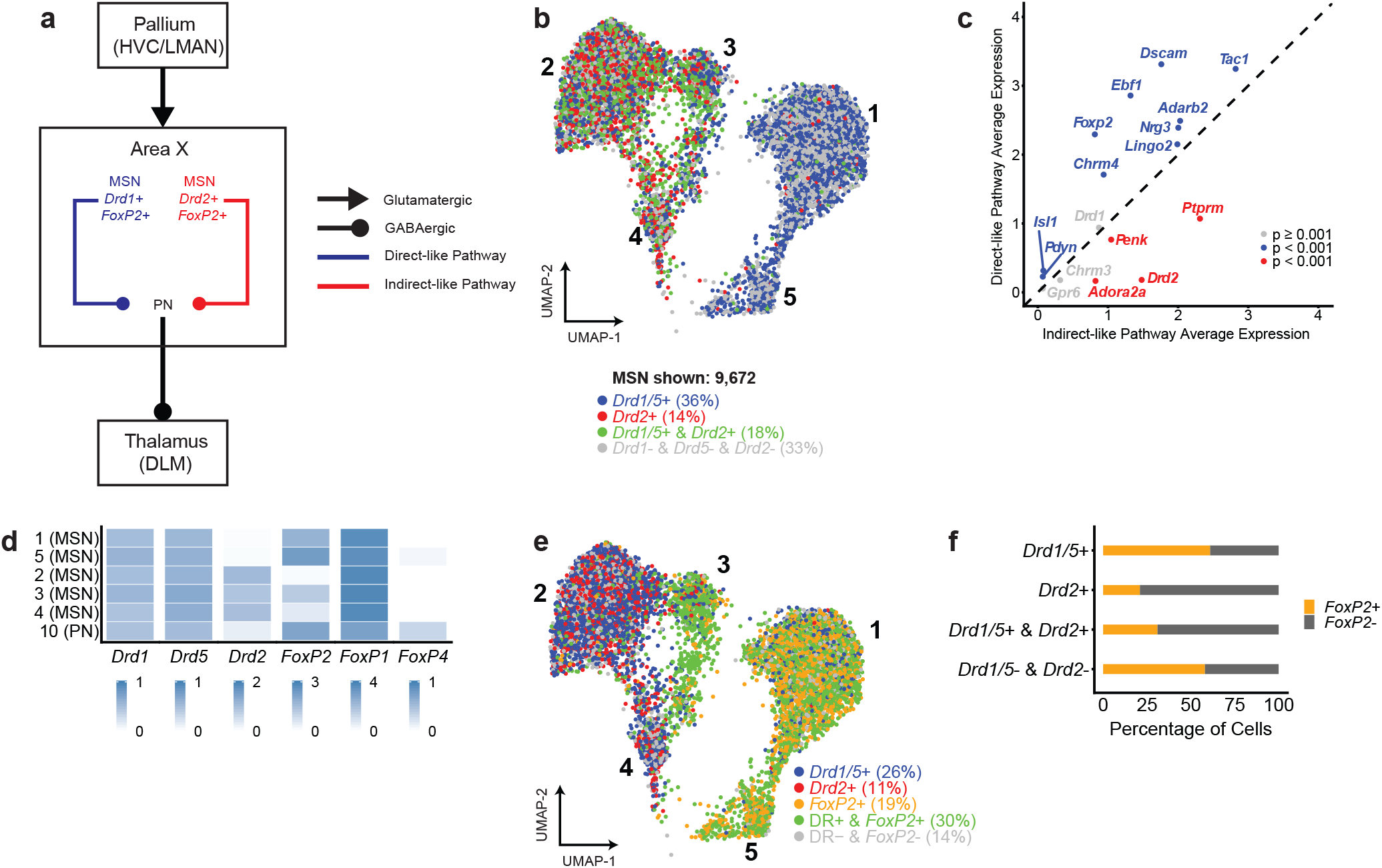
Identification of a Direct-like and Indirect-like Pathway Organization in Area X. (a) A hypothesized model for the circuitry of MSNs in Area X based on the present data. (b) UMAP projection of MSN clusters. Nuclei are colored based on the exclusive expression of *Drd1/5*, *Drd2*, both, or neither. Percentages are rounded and as a total of MSNs. (c) A differential gene expression analysis between nuclei grouped into a putative direct-like pathway (clusters 1 and 5) and indirect-like pathway (clusters 2, 3, and 4). (d) A heatmap of normalized gene expression. Expression was normalized globally across all genes, but a different scale is shown for each gene based on the highest normalized value. (e) UMAP projection of MSN clusters. Nuclei are colored based on the exclusive expression of *Drd1/5*, *Drd2*, *FoxP2*, any dopamine receptor and *FoxP2*, or none. Percentages are rounded and as a total of MSNs. (f) A stacked bar plot illustrating the percentage of *FoxP2* co-expression in nuclei classified by dopamine receptor expression.

Disruptions of FoxP2 expression have previously been shown to result in increased dopamine levels and reduced expression of dopamine receptors in the striatum ^19, 20^. Using our snRNA sequencing data we found that knockdown of FoxP2 causes an overall decrease in the *Drd1* to *Drd2* ratio across MSNs in Area X. This change in the *Drd1* to *Drd2* ratio resulted from decreased expression of *Drd1* and *Drd5* in the direct-like pathway MSNs (Figures 6a-b). Together, these results suggest that knockdown of FoxP2 alters the balance of dopaminergic receptor distributions across the direct-like and indirect-like pathways in Area X, which may drive maladaptive repetition of song syllables and/or disrupt song syntax.

**Figure 6.**
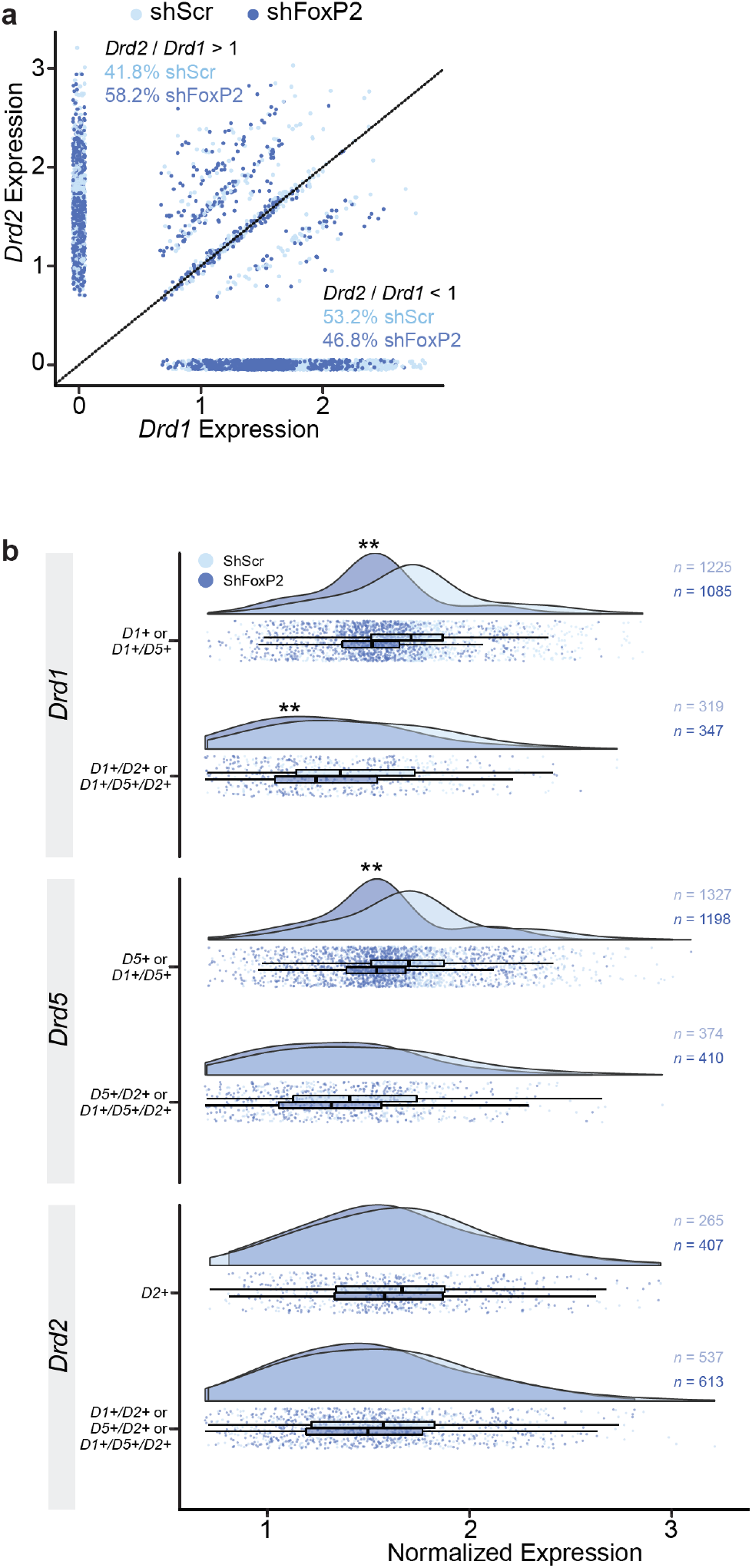
Altered Expression of Dopamine Receptors Resulting from FoxP2 Knockdown. (a) A scatterplot illustrating the correlation between *Drd1* normalized expression and *Drd2* expression in *FoxP2*+ striatal cells that are also *Drd1/5*+ or *Drd2*+. Each point represents a cell. Dashed line indicates the line of equality. Cells above the line have a ratio of *Drd2*/*Drd1* greater than 1. Cells below the line have a ratio of *Drd2*/*Drd1* below 1. (b) Distribution and boxplots of normalized expression for *Drd1*, *Drd5*, and *Drd2* in *FoxP2*+ MSNs in CS-shScr+ and CS-shFoxP2+ birds. Cells are grouped by the expression of dopamine receptors, e.g. *D1+* indicates the cells express *Drd1+* only and no other dopamine receptor, whereas *D1*+/*D5*+ indicates the cells express both *Drd1* and *Drd5*. The expression level of *Drd1* in cells from CS-shFoxP2+ birds was significantly lower than in CS-shScr+ birds (p < 0.001, Welch’s t-test) in all *Drd1*+ cells. The expression of *Drd2* in *FoxP2*+ MSNs did not differ between the two groups. The lower and upper bounds of the boxes indicate the 25th and 75th percentiles; the whiskers extend in either direction from the bound to the furthest value within 1.5 times the interquartile range.

### Phasic Dopamine in Area X Drives Maladaptive Syllable Repetitions Independent of Reinforcement Learning

Given the strong ties between FoxP2 expression and dopamine signaling^7, 19, 20, 21^, we asked if sustained manipulations of dopaminergic tone or phasic input would lead to disruptions in syllable repetition or song syntax resembling those observed following FoxP2 knockdown. Increased dopamine levels and decreased dopamine receptor expression have been associated with impairments in coordinated movements ^50, 51^, including speech. For example, childhood-onset fluency disorder, also referred to as stuttering or stammering, is associated with a hyperdopaminergic tone in the striatum ^52, 53, 54^. Dopaminergic antagonists have been shown to lessen stuttering and dopaminergic agonists are reported to transiently induce speech disturbances similar to stuttering^55, 56, 57^. We directly manipulated dopaminergic circuits across several days in freely singing birds using pharmacological or optogenetic approaches to test if and how these manipulations resulted in song disruptions and their similarity to the disruptions observed following FoxP2 knockdown.

First, we tested whether chronic elevation of dopamine could elicit gross changes in song. We implanted reverse microdialysis probes bilaterally in Area X and individually infused dopamine, Drd1-like agonist, and Drd2-like agonists, each for several days (Figures S7Sa-c). Remarkably, and in contrast to the vocal deficits observed following FoxP2 knockdown, birds continued to sing at normal rates and without disruptions in song spectral structure, syntax or syllable repetition during direct infusion of dopamine or dopamine receptor agonists (Figure S7d).

Next, we tested if sustained manipulations to phasic dopamine activity would result in disruptions of song sequences. Direct pathway MSNs, which express *FoxP2*, are thought to be sensitive to phasic increases in dopamine, while indirect pathway MSNs are thought to be more sensitive to phasic decreases in dopamine ^58^. Therefore, we systematically tested the effect of both phasic increases and phasic decreases in dopamine release over several days. Adeno-associated viruses (AAV) expressing an axon-targeted channelrhodopsin (ChR2), archaerhodopsin (ArchT), or green fluorescent protein (GFP) were injected bilaterally into VTA, and birds were implanted with optical fibers over Area X (Figures 7a-b). Bilateral optical illumination of VTA axon terminals in Area X was targeted to an individual syllable in the song motif for 3-12 consecutive days (Figures 7c and S8a). Light pulse delivery was dependent on natural trial-to-trial variation in the pitch of the targeted syllable. In agreement with our previous results using this method^59^, we found that optogenetic excitation and inhibition elicited learned changes in the pitch of the targeted syllable on future performances (Figure 7d). In addition to changes in the pitch of the targeted syllable, we found that phasic increases in dopamine also resulted in increased syllable repetition similar to those we observed in birds in which we knocked down FoxP2 expression in Area X (Figures 7e-g and S8b). Phasic excitation of VTA terminals in Area X resulted in a significant increase in the number of times birds repeated syllables at the beginning and/or end of their song motif (Figures 7f-g and S8b). This increase in syllable repetitions was observed in all ChR2 expressing birds by the third day of phasic stimulation. We did not observe disruptions in the selection and sequencing of vocal motor actions in birds that received phasic inhibition of dopamine release during singing, or in birds expressing GFP (Figures 7e and S8c-d).

**Figure 7.**
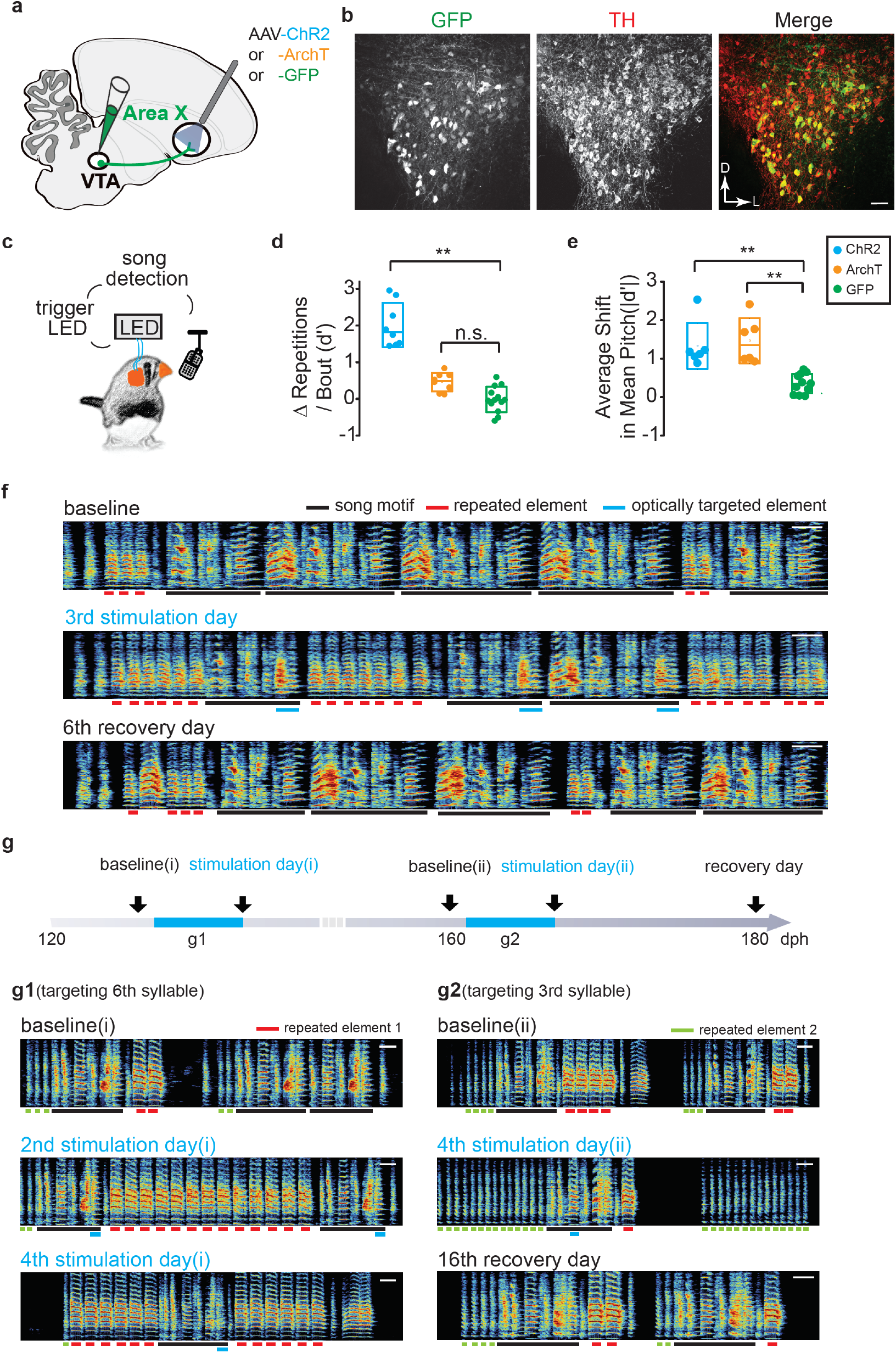
Song Contingent Excitation of Dopamine Terminals in Area X Induces Dysfluent Repetition of Song Syllables. (a) Schematic of experimental design for optogenetic manipulation of dopamine release from VTA terminals. Optogenetic construct (ChR2 or ArchT) or GFP control construct was delivered into VTA. (b) Representative coronal section through VTA shows that most neurons infected with optogenetic constructs are TH positive and located in ventral and ventrolateral portions of VTA. Scale bar, 50 μm. (c) Schematic of closed loop optogenetic experimental paradigm. (d) Changes in the number of repetitions of vocal element per song bout between the baseline day and last illumination day, expressed in units of *d*′, for ChR2+ birds (blue circles, *d*′=2.02±0.5, n=8 vocal elements from 6 birds), ArchT+ birds (orange circles, *d*′=0.46±0.21, n = 8 syllables from 6 birds), and GFP+ birds (green circles, *d*′=−0.015±0.2, n = 13 syllables from 4 birds). Change in the number of repetitions of vocal elements in ChR2+ birds were significantly greater than change in GFP+ birds (p<0.0001, Kruskal-Wallis test), but there was no significant difference between ArchT+ and GFP+ birds (p=0.12, Kruskal-Wallis test). Box indicates the median ± 1.0 SD, mean shown by open dot. (e) Average shift in mean pitch, expressed in units of |*d*′| , for ChR2+ birds (blue circles, |*d*′| =1.33±0.6, n=6 syllables from 6 birds), ArchT+ birds (orange circles, |*d*′| =1.47±0.59, n=6 syllables from 6 birds), and GFP birds (green circles, |*d*′| =0.35 , n = 11 syllables from 6 birds). Average shift in mean pitch for both ChR2+ and ArchT+ birds were higher than 0.75, and also significantly higher than control GFP+ birds (ChR2+, p=0.0025; ArchT+, p=0.0025; Kruskal-Wallis test). Box indicates the median ± 1.0 SD, mean shown by open dot. (f) Spectrograms of song recorded from a ChR2+ bird on the baseline day, 3^rd^ stimulation day and 6^th^ recovery day. Light pulses (~455 nm, 100ms) were delivered over the target syllable during lower pitch variants but not during higher pitch variants. Schematic of experiment in a ChR2+ bird in which light pulses (~455 nm, 100ms) were delivered over two different target syllables at different times over the course of two months. The bird starts repeating a song element either at the end of its motif (g1) or the beginning of its motif (g2) or both (g1 bottom), depending on which syllable in the song was optogenetically targeted. Scale bar, 200 ms.

The increased repetition of song syllables generalized to all song performances, not just optically stimulated trials, and persisted for two or more days after optical stimulations were discontinued (Figures 8d-e). This suggests that phasic increases in dopamine do not have a direct influence on ongoing vocal-motor actions but may maladaptively influence the selection and sequencing of vocal motor actions. Consistent with this interpretation, in only one case (1 of 8 birds) did the bird start to repeat the optically targeted syllable (Figure S8b). In this case the bird began to repeat a pair of syllables in the middle of its song motif, suggesting that vocal repetitions could emerge at any point in song and are not necessarily confined to the beginning and ending of the song motifs. We also found that which syllable was optically stimulated influenced which other syllable(s) birds began to repeat (Figure 7g). In a bird with a six-syllable song, optogenetic excitation of the sixth syllable (Figure 7g1) resulted in vocal repetitions at termination of the song motif, while optogenetic excitation of the third syllable (Figure 7g2) resulted in vocal repetitions at the initiation of the song motif.

Notably, optogenetic stimulation of VTA terminals did not recapitulate the full range of vocal disruptions observed following FoxP2 knockdown. We did not observe dropping of syllables or creation of new syllables following optogenetic stimulation. Together, these results suggest that FoxP2 expression in striatal circuits can influence the production and control of vocalizations in multiple ways and that relationship between phasic dopaminergic signaling and FoxP2 may more selectively disrupt precise sequencing of vocalizations and lead to maladaptive repetition of song syllables.

Since disruptions in vocal sequencing emerge while birds are also adaptively learning to change the pitch of the optically targeted song syllable, we examined if there was a relationship between adaptive (pitch learning) and maladaptive (syllable repetitions) forms of vocal plasticity. We found that optical inhibition experiments drove changes in the pitch of song syllables comparable to those seen in optical excitation experiments, yet these manipulations did not result in maladaptive vocal repetitions (Figures 7d-e). This suggests that reinforcement-based learning of changes in pitch can occur independent of maladaptive changes in syllable sequencing. In addition, we found that the recovery from optogenetic induced changes in vocal repetitions occurred on timescales that differed from recovery in pitch learning (Figures 8d-e). Recovery trajectories for these two types of learning varied from bird to bird, but their decoupling further suggests that reinforcement-based changes in how syllables are sung (pitch learning) occurs independent of maladaptive changes to circuits that may help regulate vocal sequences.

**Figure 8.**
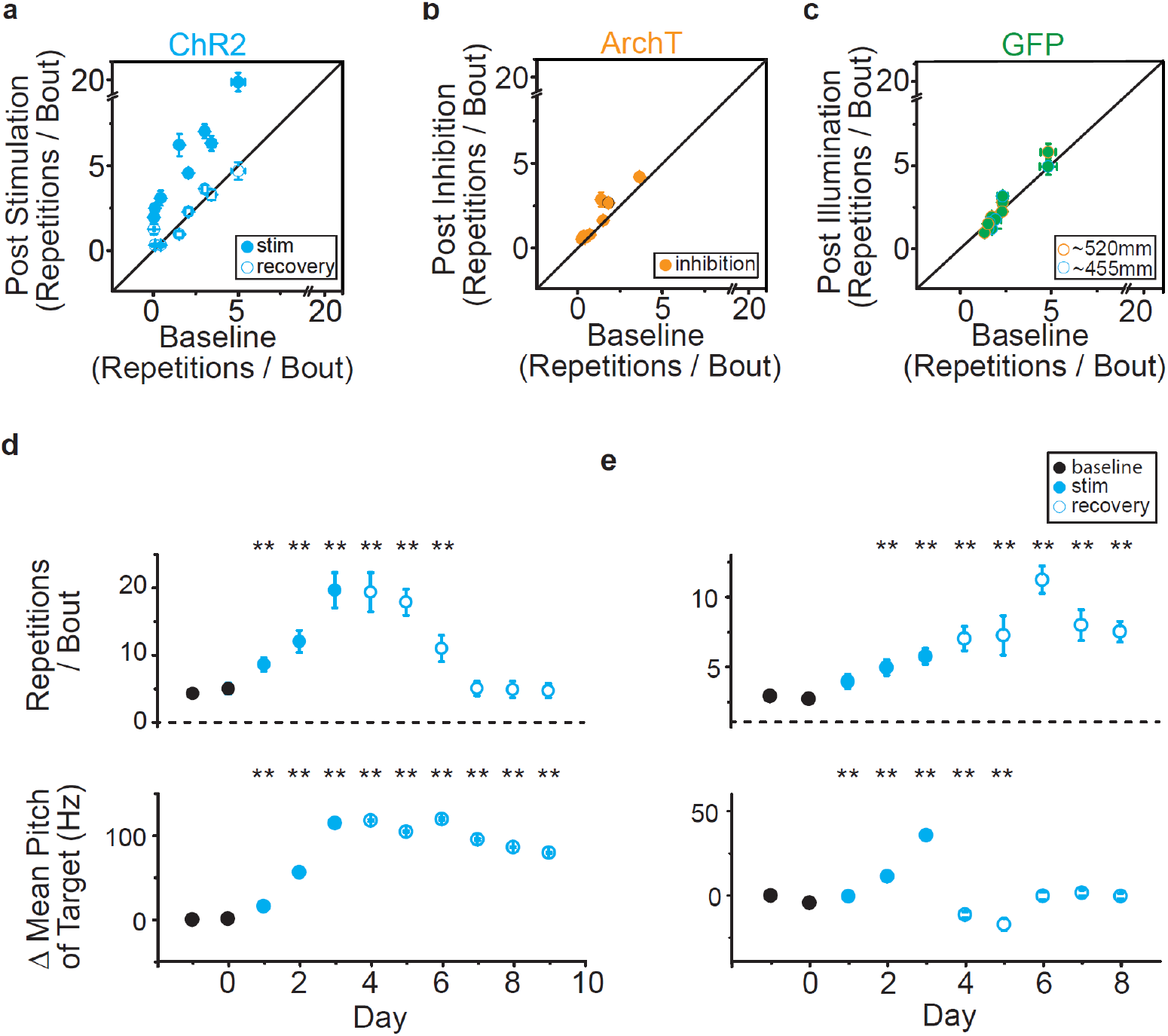
Optical Stimulation Causes Vocal Repetitions Independent of Changes in Pitch. (a) Comparison of number of repetitions of vocal element per song bout (± SEM) for the baseline day versus last stimulation day in 6 ChR2+ birds (filled, p<0.0001, Kruskal-Wallis test) or last recovery day (open, p>0.35, Kruskal-Wallis test, n=8 vocal elements). The number of repetitions of vocal elements in the last stimulation day was significantly higher than the number of repetitions in either the baseline day or the last recovery day (p=0.0081, Friedman test), but not between the baseline day and the last recovery day (p>0.99, Friedman test). (b) Comparison of number of repetitions of vocal element per song bout (± SEM) for the baseline day versus last inhibition day in 6 ArchT+ birds (filled, p>0.08; filled with black outline p=0.028, Kruskal-Wallis test, n=8 syllables). (c) Same as (b), but for GFP+ birds (blue outline, p>0.8, Kruskal-Wallis test, n=6 syllables from 3 birds; orange outline, p>0.15, Kruskal-Wallis test, n=7 syllables from 3 birds). Blue and orange outline indicate birds illuminated by LED with wavelength of ~455 nm or 520 nm, respectively. (d) The number of repetitions of a syllable from bird shown in Fig. 7f per song bout (top) and changes of mean pitch(bottom) of the optically targeted syllable in Fig. 7f) during baseline (black circles, day −1 and 0), stimulation (blue circles, day 1-3), and recovery days (open blue circles, day 4-9). The number of repetitions of the affected syllable (red line in Fig. 5D) per song bout was significantly increased during day 1-6 relative to baseline (p<0.01, Kruskal-Wallis test), whereas changes in the mean pitch of the target syllable during day 1- 9 were significantly higher than changes in the baseline day (p<0.01, Kruskal-Wallis test). (e) same as (d), but for another bird. The number of repetitions of the affected syllable per song bout was significantly increased during day 2-8 relative to baseline (p<0.01, Kruskal-Wallis test), whereas changes in the mean pitch of the target syllable during day 1- 5 were significantly higher than changes relative to baseline (p<0.01, Kruskal-Wallis test).

Together, these findings indicate that increased phasic excitation of dopaminergic VTA terminals in Area X causes problems with initiating and terminating song, marked by birds perseverating on syllables at the beginning and ending of song. That only phasic excitation is sufficient to drive these disruptions raises the possibility that the direct pathway through the striatum, and FoxP2 expressing neurons in this circuit, functions in fluently starting and stoppings song bouts in zebra finches.

## DISCUSSION

Precise sequencing of vocal motor actions is necessary for vocal communication. While recent studies have begun to clarify the role of FoxP2 and reinforcement signals in learning how to properly produce individual syllables ^22, 23, 59, 60, 61, 62^, the identity of cells and circuits that control selection and sequencing of syllables have remained unclear. We show that the expression of FoxP2 is critical for the maintenance of adult vocalizations and that knockdown of its expression causes syllable repetitions and disruptions to song syntax. Restoring FoxP2 expression later in adulthood resulted in recovery of linear song syntax. This adult plasticity is particularly striking given that birds of this age are thought to have limited song plasticity and be less reliant on auditory feedback to maintain their songs ^29^. We identify phasic dopamine as a selective mediator of maladaptive changes in song sequences and therefore show that dopamine not only functions in reinforcement-based learning of song syllables^59^ but also in the accurate sequencing of syllables in adulthood.

We identified distinct populations of *Drd1*+/*FoxP2*+ MSNs and *Drd2*+/*FoxP2*− MSNs in Area X, consistent with striatal direct and indirect MSN populations seen across other vertebrates^45^. Nearly 20% of the MSNs in Area X co-express *Drd1* and *Drd2*. Although this proportion is less than what has been previously reported, with most Area X MSNs thought to co-express both genes ^47, 48^, it is much higher than what has been reported in the mammalian striatum, where MSNs co-expressing *Drd1* and *Drd2* only represent about 1-4% of total MSNs ^32, 44, 46^. The broad groupings of MSNs into direct-like and indirect-like pathways identified here highlight commonalities in the gene markers between Area X and mammalian pathways ^44, 46, 49^; however, this is not the case for all genes. While *Drd2* and *FoxP2* are differentially expressed in MSN neuronal clusters, *Drd1* is not. Furthermore, a substantial portion (33%) of the MSNs in our data set do not express any dopamine receptor. Although this may reflect the inability to detect transcripts expressed at low levels, 33% is much higher than what has been reported in mammalian striatum ^32, 44, 46^.

Heterozygous mutations of FOXP2 cause Childhood Apraxia of Speech, also referred to as Developmental Verbal Dyspraxia. This speech impairment is thought to result in part from disruptions in developmental plasticity of basal ganglia circuits ^63^. The role of FOXP2 in the maintenance of adult speech is not known. Altering the expression of FoxP2 in Area X impairs song imitation in juvenile birds^16^, but it had not been shown to be necessary for normal maintenance of the adult song. A previous study of FoxP2 knockdown in adult Area X focused on changes in context dependent syllable variability^19^. Using the same hairpin sequence used in previous studies ^16, 19^, but expressing it in Area X using a novel AAV CS-shFoxP2 construct rather than lentiviral based methods, we show that FoxP2 expression is necessary for maintenance of adult song sequences and syntax. Although we did not observe changes in syllable variability following FoxP2 knockdown, we did not measure social context dependent singing in this study. We made this choice because female directed singing has been shown to elevate dopamine in Area X as well as increase the number of introductory notes birds sing^17, 64, 65, 66^, variables that we wished to manipulate and quantify, respectively, in the undirected singing condition. We suspect that the broader neural expression in Area X afforded by AAV over lentivirus could account for the stronger effects on song performance and maintenance reported here^67^. Knocking down FoxP2 expression in Area X of adult zebra finches drove a significant increase in the repetition of song syllables, the elimination of certain syllables from the song, and the improvisation of new syllables. However, we did not observe spectral degradation of individual song syllables, as seen following deafening^29, 30, 68^. This indicates that the ability to control fine-scale features of song was largely undisturbed, while global control of selection and sequencing of syllables was impaired.

When we rescued FoxP2 knockdown in adult birds, we found that they were able to recover normal song syntax within approximately 3 months. This suggests that the progressive and maladaptive changes in behavior driven by disruptions of FoxP2 can in part be overcome with restoration of gene expression. Despite its name, Childhood Apraxia of Speech is a lifelong condition. Finding that birds can recover normal song syntax well into adulthood, a timepoint when song behavior is thought to be mostly rote ^29^, suggests the possibility that genetic therapies for speech disorders could have relevance even beyond early developmental windows when speech is first learned.

We demonstrate that FoxP2 knockdown and phasic activation of VTA terminals in Area X similarly disrupt the fluent initiation and termination of adult song by increasing the repetition of song syllables. This is consistent with the excess dopamine hypothesis of stuttering, which propose that a hyperdopaminergic tone in the basal ganglia causes speech dysfluencies, including maladaptive vocal repetitions and difficulty initiating speech ^52, 53, 54^. Elevated dopamine levels in the striatum are reported in association with disruptions of FoxP2 expression and with imbalanced expression of dopamine receptor in direct and indirect pathways^7, 69^. Although we did not directly measure dopamine levels in Area X, we reason that the decreased expression of *Drd1/5* in the direct-like pathway MSNs is compensated for by elevation of dopamine levels in the striatum, as has been previously reported in mammals^7, 20^. Further, we only observed disruptions in singing following song-contingent phasic activation of VTA terminals, and not during chronic infusion of dopamine or dopamine receptor agonists. This suggest a strong behavior or vocal sequence specific contingency in how elevated dopamine influences vocal fluency. It is common in stuttering to have particular words or sounds that different individuals find particularly problematic to produce. Understanding if or how dopamine is dynamically regulated during production of these and other less problematic words or sounds may be particularly revealing to the underlying circuit activity disruptions associated the stuttering.

When considered within the context of the broader literature, it appears that Area X plays a dual role in song. It is involved in learning how individual song syllables should be sung and in controlling larger scale selection and sequencing of these syllables^11, 12, 13, 24, 25, 26^. Positive and negative dopaminergic reinforcement signals guide how individual song syllables are sung on future performances^23, 59^, while disruptions to the contributions of the direct-like and indirect-like pathways may regulate syllable selection and sequencing (current results). Knockdown of FoxP2 leads to a decrease in *Drd1*/*Drd2* ratios, which may drive an imbalance in direct-like and indirect-like pathways. Continued phasic excitation of dopaminergic inputs, which may preferentially influence activity in the direct pathway ^58^, consistently resulted in birds having prominent sequence disruptions/repetitions at the beginning and end of their song. Similarly, expression of the mutant gene fragment that causes Huntington’s disease in Area X also causes disruptions in song syllable selection and repetition ^24^. These disruptions, however, tend to be more restricted to changes in core aspects of the song and do not accumulate at the initiation and termination of song. Indirect pathway neurons are particularly vulnerable at early stages of Huntington’s disease ^51, 70, 71, 72^, which together with our findings, raises speculation that disruptions in the direct-like pathway could more readily cause vocal repetitions at initiation and termination of vocal-motor sequences, whereas disruptions in the indirect-like pathway could tend to disrupt sequences in the middle of song. Hierarchical representations of song sequences may therefore rely critically on coordinated activity of the direct-like and indirect-like pathways in Area X and the precise timing signals that facilitate transitions between individual syllables. The molecular cataloging of Area X cell types, and the tools for reversible genetic manipulations described here, provide the means to start testing these and related hypotheses about the selection and sequencing of vocal-motor actions.

Many speech disorders arise from problems in translating volitional speech plans into accurate motor actions ^2, 73, 74, 75^ and have been linked to hyperdopaminergic signaling in the striatum ^52, 53, 54, 76^. Together, our findings in Area X indicate that convergent circuit mechanisms may be involved in translating volitional birdsong and speech plans into to fluent vocal-motor actions.

## MATERIALS AND METHODS

### Animals

All experiments were performed on adult male zebra finches (*Taeniopygia guttata*) raised in a breeding facility at UT Southwestern and housed with their parents until at least 50 days of age. During experiments, birds were housed individually in sound-attenuating recording chambers (Med associates) on a 12/12 h day/night cycle and were given ad libitum access to food and water. All procedures were performed in accordance with established protocols approved by the UT Southwestern Medical Center Animal Care and Use Committee.

### Plasmid Construction and Viral Vectors

The backbone of CS constructs was based on pAAV-EF1α-DO-mCherry (Addgene, #37119)^27^, and the fluorescent protein cDNA for tagBFP was cloned from pdCas9::BFP-humanized (Addgene, #44247). A zebra finch FoxP2 cDNA clone provided by Erich Jarvis was subcloned into pLenti6.4 using a gateway reaction, adding a V5 tag. Two hairpins (shFoxP2a, target sequence AACAGGAAGCCCAACGTTAG T^77^, and shFoxP2i, target sequence ACTCATCATTCCATAGTGAAT) were inserted downstream of the U6 promoter at the base of the Mir-30 stem-loop. The scrambled hairpin (shScr, sequence CCACTGTACTATCTATAACAT) was designed as a control. Hairpins were then assembled into the pTripZ vector (Thermo Scientific, MA, USA) by directional ligation into the XhoI-EcoRI cloning sites. We then replaced the EF1α promoter with a CAG promoter and assembled hairpins together with pTripZ context sequence (between the BspD1-MluI cloning sites) in the forward orientation and tagBFP transgene in the reversed orientation downstream of the mCherry transgene. All N Terminal sites included a Kozak sequence (GCCACC) directly preceding the start codon. Sequence confirmation was done by the McDermott Center Sequencing Core at UT Southwestern Medical Center. The recombinant AAV vectors were amplified by recombination deficient bacteria, One Shot Stbl3(C737303, Invitrogen, CA, USA), serotyped with AAV1 coat proteins and produced by the University of North Carolina vector core facility (Chapel Hill, NC, USA) with titer exceeding 10^12^ vg/ml, the Duke viral vector core facility (Durham, NC, USA), IDDRC Neuroconnectivity Core in Baylor College of Medicine (Huston, TX, USA ) or in the Roberts lab with titer exceeding 10^11^ vg/ml. All viral vectors were aliquoted and stored at –80 °C until use.

### Stereotaxic Surgery

All surgical procedures were performed under aseptic conditions. Birds were anesthetized using isoflurane inhalation (1.5-2%) and placed in a stereotaxic apparatus. Viral injections and cannula or microdialysis probe implantation were performed using previously described procedures^59^ at the following approximate stereotaxic coordinates relative to interaural zero and the brain surface were (rostral, lateral, depth, in mm): Ov (2.8, 1.0, 4.75), the center of Ov was located and mapped based on its robust white noise responses; VTA relative to the center of Ov (+0.3, −0.2, +1.8); Area X (5.1, 1.6, 3.3) with 43-degree head angle or (5.7, 1.6, 3) with 20-degree head angle, the boundary of Area X was verified using extracellular electrophysiological recordings. 0.7-2 μl AAVs were injected according to the titer of constructs and allowed 3-8 weeks for expression before birds were subjected for behavioral tests, immunohistochemistry and/or sequencing experiments.

### Behavioral Assays

#### Song Recording

Acoustic signals were recorded continuously by a microphone immediately adjacent to the bird’s cage using Sound Analysis Pro2011^78^ and bandpass filtered between 0.3 and 10 kHz. All songs presented were recorded when the male was isolated in a sound-attenuating chamber.

#### Optogenetic Manipulation of VTA Axon Terminals in Behaving Birds

All procedures were reported previously ^59^. Briefly, male birds were randomly assigned bilateral injection of either AAV-ChR2, -ArchT or GFP constructs in VTA at ~70 dph. Birds were implanted with fiber optics after they were at least ~100 dph. Birds were given at least 1 week to recover from cannula implantation and to habituate to sing with attached optical fibers. Custom LabView software (National Instruments) was used for online detection of predetermined target syllables and implementation of closed-loop optogenetic manipulation^79^. 100-ms light pulses were delivered over a subset of variants of the target syllables in real time (system delay less than 25ms) for 3-12 consecutive days, as described previously^59^. Investigators were not blinded to allocation of optogenetic experiments. 3-5 mw of ~455nm and 1.5-4 mw of ~520nm LED output was delivered from the tip of the probe (200 or 250um, NA=0.66, Prizmatix, Israel) to ChR2+ and ArchT+ birds, respectively, while either ~455nm or ~520nm light pulses were delivered to GFP+ birds. ChR2+(n=6) and ArchT+ (n=6) birds with significant changes in the pitch of target syllables (|*d*′| > 0.75 significance threshold) were included for subsequent behavioral analyses.

#### Pharmacological Manipulation of DA Circuit in Behaving Birds

We used two microdialysis systems to chronically infuse dopamine hydrochloride (DA), SKF 38393 hydrobromide (SKF) or (-)-Quinpirole hydrochloride (Qui) (#3548, #0922 and #106, Tocris, MN, USA) into Area X to simulate tonically elevated DA levels. Two male adult birds were implanted bilaterally with probes constructed in house from plastic tubing (427405, BD Intramedic, PA, USA; 27223 and 30006, MicroLumen, FL, USA) which served as a drug reservoir, fitted at the end with a 0.7mm-long semipermeable membrane (132294, Spectra/Por, MA, USA) allowing drug to slowly diffuse into the brain throughout the day^80, 81^. Freshly made DA, SKF or Qui(400mM) were used to fill and refilled microdialysis probes every morning for 4 consecutive days following 3 days of dialysis with PBS. Two other birds were implanted with guide cannula (8010684, CMA, MA, USA) bilaterally over Area X and microdialysis probes (1mm membrane length, 6kDa cutoff, P000082, CMA, MA, USA) were not inserted until birds recovered from surgery and were singing (2-3 days) as described previously^82, 83^. Fresh made DA, SKF or Qui(100-400mM) or PBS (for baseline) were continuously delivered to Area X for 3 – 8 consecutive days at a rate of 0.2 μl/min via a fluid commutator connected to a syringe pump outside the bird’s isolation chamber. In all cases birds could comfortably move and sing during infusion.

### Behavioral Analysis

#### Song structure

Zebra finch song can be classified into three levels of organization: syllables, which are individual song elements separated by short silent gaps >5 ms in duration; motifs, which are stereotyped sequences of syllables (outlined by black lines throughout the manuscript); and song bouts, which are defined as periods of singing comprised of introductory elements (eg. syllables indicated by red lines in Figure 2b), followed by one or more repeats of the song motif with inter-motif intervals >500 ms^65, 84^.

#### Quantification of vocal repetition

The vocal element being repeated consisted of either a single syllable (e.g. introductory element repeated in Figure 2b or ending song syllable repeated in Figure 2c) or multiple syllables (e.g. song syllables repeated in Figure S8b). The number of repetitions of individual vocal element per song bout(n) is defined as the total number of consecutively repeated vocal elements, not including the first rendition each time the element is sequentially produced within the song bout. To calculate the number of motifs per bout(m), we counted all motifs in which at least the first half of the motif was produced within a song bout. The number of repetitions of individual vocal elements per motif is defined as n/m. *d*′scores were computed to express the changes in the mean number of repetitions of individual vocal element per song bout(n) relative to the last baseline day^59^:

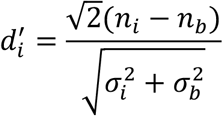

*n*_*i*_ is the mean number of repetitions of individual vocal element per song bout on day *i* and 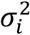 is the variance on day *i*. Subscript b refers to last baseline day. In the case of equal variances 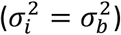, 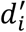 reports the changes in average repeats per bout between training day *i* and the baseline day in the convenient unit of SDs. Zebra finch song is mostly linear, exhibiting few repetitions. However, birds do tend to repeat introductory elements and ending elements a small number of times. We focused analysis on vocal elements or song syllables that were observed to repeat two or more times on at least one occasion during optical manipulations or following knockdown of FoxP2 in Area X. Once identified, we retrospectively tracked these syllables back to baseline periods or forward in time to recovery periods in order to examine if repetitions changed during our manipulations. Syllables which were not repeated during optical manipulations or following knockdown of FoxP2 were not further considered.

#### Quantification of changes in sequencing

The frequency of syntax altering is defined as the chance of new syllable transition(s), which was not identified before FoxP2 knockdown, occurring in a song bout. The frequency of omitting syllables is defined as the chance of pre-existing syllable(s), which was identified before FoxP2 knockdown, being absent in a song bout. The frequency of adding syllables is defined as the chance of de novo created syllable(s), which was not identified before FoxP2 knockdown, being present in a song bout. Omitting or adding syllables in the beginning or end of a motif identified before FoxP2 knockdown, regardless whether new syllable transition(s) occur(s) as a result, is excluded from being classified as syntax altering.

### Immunohistochemistry and Immunoblotting

Birds were anesthetized with Euthasol (Virbac, TX, USA) and transcardially perfused with ice cold phosphate buffered saline (PBS), followed by 4% paraformaldehyde (PFA) in PBS. The brains were post-fixed in 4% PFA for 2hrs at 4 °C, then transferred to PBS containing 0.05% sodium azide. The brains were sectioned at 50 μm, using a Leica VT1000S vibratome. Immunohistochemistry (IHC) was performed as described previously^59^. Sections were first washed in PBS, then blocked in 10% normal donkey serum (NDS) in PBST (0.3% Triton X-100 in PBS) for 1hr at room temperature (RT). Sections were incubated with primary antibodies in blocking solution (PBST with 2% NDS and 0.05% sodium azide) for 24-48 hrs at 4 °C, then washed in PBS before a secondary antibody incubation (703-545-155, 711-475-152, 705-605-003, 711-585-150 or 711-585-152, Jackson Immuno Research, ME, USA) for 4-6 hrs at RT. Images were acquired with an LSM 710 laser-scanning confocal microscope (Carl Zeiss, Germany), processed in Zen Black 2012 and analyzed in ImageJ. To minimize bias during quantification, matched regions across animals were selected for IHC according to GFP expression levels. To mitigate any cross-talk between distinct fluorescent channels, spectral unmixing was used and a regions of interest (ROI) around each mCherry+ or tagBFP+ cell was manual drawn and overlaid to the channel (Alexa Fluor 647) designed for FoxP2. The expression level of FoxP2 in each ROI was estimated by measuring the mean intensity value from the nucleus of each ROI and subtracting the background intensity.

The expression level of FoxP2 in control cells are estimated from 20 random non-infected cells (FoxP2+mCherry-tagBFP-) from the same slice where the expression in the mCherry+ and tagBFP+ cells are measured.

Immunoblotting (IB) was carried out as described previously^85^. Cell lysates from each sample were separated by SDS-PAGE and transferred to an Immuno-Blot PVDF Membrane (162-0177, Bio-Rad Lab., CA, USA), then blocked with 1% skim milk in TBST (tris-buffered saline with 0.1% Tween-20) for 1 hr at RT. The membrane was incubated with primary antibodies overnight at 4°C, washed with TBST, and reacted with the appropriate horseradish peroxidase (HRP)-conjugated species-specific secondary antibodies (NA934 and NA931, Sigma-Aldrich, MO, USA; AP180P, Millipore, MA, USA) for 1 hr at RT. The signals were detected by Clarity western ECL substrate (170-5060, Bio-Rad Lab., CA, USA).

The primary antibodies used were: Goat anti-FOXP2(ab1307, Abcam, MA, USA), Goat anti-FOXP2(sc-21069, Santa Cruz Bio., TX, USA), mouse anti V5 tag(R960-25, Invitrogen, CA, USA), rabbit anti-RFP (mCherry, 600-401-379, Rockland, PA, USA), mouse anti-RFP (mCherry, 200-301-379, Rockland, PA, USA), rabbit anti-GFP rabbit (A11122, Invitrogen, CA, USA), chicken anti-GFP (AB16901, Millipore, MA, USA), rabbit anti-tRFP (tagBFP, AB233, Evrogen, Moscow, Russia) and mouse anti-GAPDH (MAB374, Millipore, MA, USA). The specificity of primary antibodies against FoxP2, RFP or GFP were confirmed by two independent primary antibodies for both IHC and IB.

### Statistics

Shapiro-Wilk Test was performed for all behavioral data to test for normality of underlying distributions. Unless otherwise noted, statistical significance was tested with non-parametric statistical tests; Wilcoxon signed-rank tests and Wilcoxon rank-sum tests were used where appropriate. Kruskal-Wallis test was performed when comparisons were made across more than two conditions (e.g., baseline vs stimulation day vs recovery day) from individual animal, whereas Friedman tests were performed when data was pooled across animals and comparisons were made across more than two conditions. Statistical significance refers to *p < 0.05, **p < 0.02. Statistical details for all experiments are included in their corresponding figure legends.

### Tissue Processing for single-nucleus RNA Sequencing (snRNA-seq)

Adult zebra finches (120-140dph) were injected with CS-shFoxP2(n=2) and CS-shScr(n=2) constructs and scarified at 180-200dph. Birds were put down prior to lights in the to ensure that singing behavior did not affect our results. Each bird was rapidly decapitated, and its brain was placed in ice-cold ACSF (126 mM NaCl, 3 mM KCl, 1.25 mM NaH_2_PO_4_, 26 mM NaHCO_3_, 10 mM D-(+)-glucose, 2 mM MgSO_4_, 2 mM CaCl_2_) bubbled with carbogen gas (95% O2, 5% CO2). The cerebellum was removed with a razor blade and the cerebrum was glued to a specimen tube for sectioning with a VF-200 Compresstome (Precisionary Instruments). Coronal 500 μm sections were made in ice-cold ACSF and allowed to recover in room temperature ACSF for 5 min. Area X punches were placed into a tube containing ACSF on ice until all punches were collected and pooled from 2 birds per condition. Tissue punches were dounce-homogenized in 500 μl ice-cold Lysis Buffer (10 mM Tris pH 7.4, 10 mM NaCl, 3 mM MgCl_2_, 0.1% IGEPAL CA-630) and transferred to a clean 2 ml tube. Then, 900μl of 1.8 M Sucrose Cushion Solution (NUC201-1KT, Sigma, MO, USA) was added and pipette-mixed with nuclei 10 times. 500 μl of 1.8 M Sucrose Cushion Solution was added to a second clean 2 ml tube, and the nuclei sample was layered on top of the cushion without mixing. The sample was centrifuged at 13,000 × g for 45 min at 4°C and all but ^~^100 μl of supernatant was discarded to preserve the pellet. The pellet was washed in 300 μl Nuclei Suspension Buffer (NSB) (1% UltraPure BSA (AM2618, Thermo Fisher Scientific, MA, USA) and 0.2% RNase inhibitors in PBS) and centrifuged at 550 × g for 5 min at 4°C. All but ~50 μl of supernatant was discarded and the pellet was resuspended in the remaining liquid and filtered through a FLOWMI 40 μm tip strainer (H13680-0040, Bel-Art, NJ, USA). Samples were diluted to 1000 nuclei/μl with NSB for targeting 10,000 nuclei for snRNA-seq. Libraries were prepared using the Chromium Single Cell 3’ Library & Gel Bead Kit v3 according to the manufacturer’s instructions and sequenced using an Illumina NovaSeq 6000 at the North Texas Genome Center at UT Arlington.

### Pre-processing of snRNA-seq Data

Raw sequencing data was obtained as binary base cells (BCL files) from the sequencing core. 10X Genomics CellRanger v.3.0.2 was used to demultiplex the BCL files using the *mkfastq* command. Extracted FASTQ files were quality checked using FASTQC v0.11.5. Paired-end FASTQ files (26 bp long R1 – cell barcode and UMI sequence; 124 bp long R2 – transcript sequence) were then aligned to a reference Zebra Finch genome (bTaeGut1_v1.p) from UCSC Genome Browser, and reads were counted as number of unique molecular identifiers (UMIs) per gene per cell using 10X Genomics CellRanger v.3.0.2 *count* command. Since the libraries generated are single-nuclei libraries, reported UMIs per gene per cell accounts for reads aligned to both exons and introns. This was achieved by creating a reference genome and annotation index for pre-mRNAs.

### snRNA-seq Clustering Analysis

The resulting count matrices from the data pre-processing steps were analyzed with the Seurat analysis pipeline in R (v.3.0, https://satijalab.org/seurat/v3.0/pbmc3k_tutorial.html). Cells with more than 10,000 UMI and more than 5% mitochondrial genes were filtered out to exclude potential doublets and dead or degraded cells (Figures S5c-f). As described in the Seurat pipeline, the data were log-normalized and scaled using a factor of 10,000 and regressed to the covariates of the number of UMI and percent mitochondrial genes. Top variable genes were identified and principal components (PCs) were calculated from the data. PCs to include were identified by “ElbowPlot” in Seurat, where PCs are ranked according to the percentage of variance each one explains; PCs were excluded after the last noticeable drop in explanatory power. With the selected PCs, the Louvain algorithm was then used to identify clusters within the data. Clusters were visualized with uniform manifold approximation and projection (UMAP) in two dimensions. For hierarchical clustering, differentially expressed genes across the dataset were identified with the Wilcoxon Rank Sum test (Seurat FindAllClusters, min.pct = 0.1, logfc.threshold = 0.25, max.cells.per.ident = 200), the normalized data was averaged within each cluster, and the 50 genes with the top fold-change for each cluster were then used for hierarchical clustering (function pvclust from the R package *pvclust v2.2-0*, correlation distance, average agglomeration, 100 bootstrap replicates).

### snRNA-seq Dataset Integration and Differential Gene Expression

In order to compare gene expression between CS-shScr+ and CS-shFoxP2+ birds, the datasets must first be combined. Each dataset was processed independently as described in Clustering Analysis up until the point of normalization. Once normalized, the datasets were integrated as described in the Seurat integrate pipeline. The integrated dataset was then regressed to the covariates of the number of UMI and percent mitochondrial genes. The subsequent analysis proceeded as described in Clustering Analysis. The integrated data were used to generate the integrated clusters and UMAP plot, as recommended by the Seurat pipeline (https://satijalab.org/seurat/v3.0/immune_alignment.html). Comparisons of gene expression across clusters, and across datasets, were pulled from each individual RNA assay, as recommended by the Seurat pipeline. Differentially expressed genes were identified using the Wilcoxon ranked sum test with the Bonferroni correction as implemented in the Seurat pipeline.

### Data and Code Availability

The NCBI Gene Expression Omnibus (GEO) accession number for the snRNA-sequencing data in this manuscript is GSE136086. Codes for data pre-processing, clustering, and differential gene expression analysis are available on GitHub (https://github.com/konopkalab/songbird_areax).

### Cluster Function Assignment

Clusters were assigned functional identities based on the expression of gene markers established in the literature (see Table 1 below). Identity names were as specific as possible given the confidence and clarity of the gene expression pattern in relation to expression in other clusters. When gene markers have been established in Area X in zebra finches specifically in addition to mammals, this is noted in the table with a “ZF”. For downstream analyses, specific UMAP coordinates were used to further specify MSNs (UMAP_1 > −5 & UMAP_2 > −6) and PNs (UMAP_1 > 0 & UMAP_2 > −7.5).

**Table 1.**

| Gene Symbol | Gene Name | Identity | Reference |
| --- | --- | --- | --- |
| *Gad2* | Glutamate decarboxylase 2 | GABAergic | <sup>32</sup> |
| *Slc17a6* | Solute carrier family 17 member 6 | Glutamatergic | <sup>32</sup> |
| *Lhx6* | LIM homeobox 6 | MGE-derived | <sup>86</sup> |
| *Pvalb* | Parvalbumin | Interneuron (ZF) | <sup>31</sup> |
| *Sst* | Somatostatin | Interneuron (ZF) | <sup>31</sup> |
| *Npy* | Neuropeptide Y | Interneuron (ZF) | <sup>31</sup> |
| *Nos1* | Nitric oxide synthase 1 | Interneuron (ZF) | <sup>31</sup> |
| *Chat* | Choline O-acetyltransferase | Interneuron (ZF) | <sup>31, 37</sup> |
| *Penk* | Proenkephalin | PN (ZF) | <sup>37</sup> |
| *Tshz1* | Teashirt zinc finger homeobox 1 | PN (ZF) | Present study |
| *Lrig1* | Leucine rich repeats and immunoglobulin like domains 1 | Astrocyte | <sup>87</sup> |
| *Meis2* | Meis homeobox 2 | LGE-derived | <sup>86</sup> |
| *Ppp1r1b* | Protein phosphatase 1 regulatory subunit 1B (also known as DARPP-32) | MSN (ZF) | <sup>31, 32</sup> |
| *FoxP1* | Forkhead box P1 | MSN (ZF) | <sup>33</sup> |
| *FoxP2* | Forkhead box P2 | MSN (ZF) | <sup>15, 33</sup> |
| *Tac1* | Tachykinin precursor 1 | MSN (ZF) | <sup>31, 37</sup> |
| *Csf1r* | Colony stimulating factor 1 receptor | Microglia | <sup>87</sup> |
| *Flt1* | Vascular endothelial growth factor receptor 1 | Endothelial | <sup>87</sup> |
| *Mbp* | Myelin basic protein | Oligodendrocyte | <sup>87</sup> |
| *Pdgfra* | Platelet-derived growth factor receptor A | Oligodendrocyte Precursor | <sup>87</sup> |

## Acknowledgments

The authors thank members of the Roberts and Konopka laboratories for discussion and comments on the manuscript, Jennifer Holdway and Matthew Harper for laboratory support, Gaurav Chattree for helping with optogenetic experiments, Maaya Ikeda and Chung Yan Cheung for advice on reverse microdialysis experiments, Andrea Guerrero for recording of directed singing behavior, Ashley Anderson for cloning of FoxP2 V5, and Erich Jarvis for the original clone of zebra finch FoxP2.

## Author contributions

LX, DPM, and TFR designed the experiments and wrote the manuscript, LX collected and analyzed the optogenetic, pharmacological, and gene knockdown experiments, and help collect the snRNA-seq data, DPM analyzed the snRNA-seq data, MC^2^ analyzed and imaged the anatomical data and helped interpret the directed singing data, MC^1^ collected the snRNAseq data, AK developed bioinformatic pipeline for snRNA-seq data analysis and helped analyze the data, GK supervised the snRNA-seq data collection and analysis and helped design the reversible gene knockdown experiments, TFR supervised all experiments. All authors read and commented on the manuscript.

## Data and materials availability

All data is available in the main text or the supplementary materials.

## SUPPLEMENTAL INFORMATION

**Figure S1.**
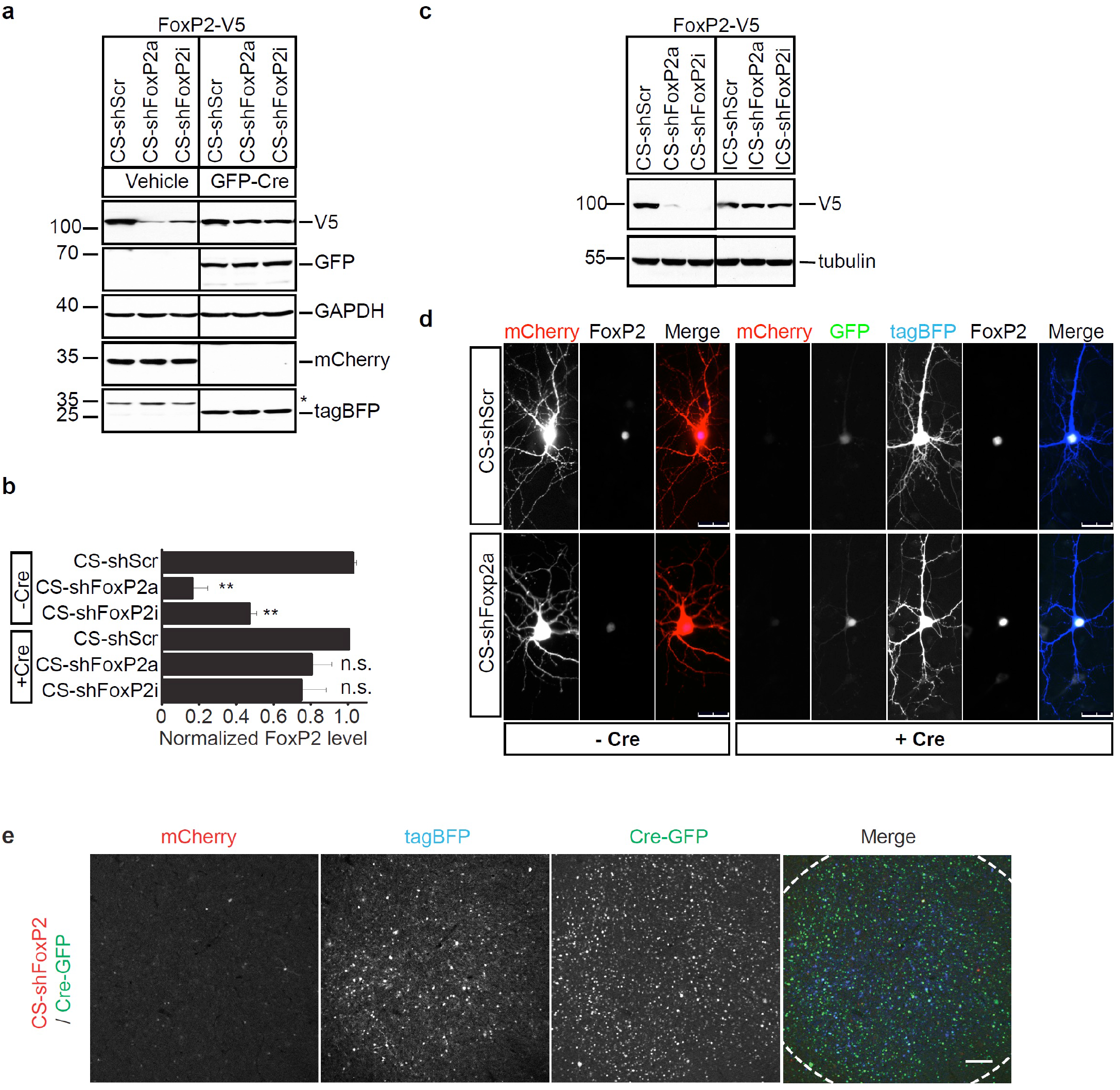
Validation of CS constructs *in vitro* and *in vivo*. (a) Validation of CS constructs in vitro with western blotting. Two independent small hairpin RNAs against zebra finch FoxP2 gene (CS-shFoxP2a & i) and scrambled hairpin (CS-shScr) together with V5-tagged zebra finch FoxP2 were co-transfected into HEK stable cell lines expressing either vehicle or GFP-Cre as indicated. At 72 h after transfection, lysates of cells were subjected to immunoblotting with V5(to detect FoxP2), GFP (to detect Cre-GFP), GAPDH, RFP (to detect mCherry) and tagBFP antibody. (b) The expression levels of FoxP2 were quantified from three independent experiments shown in (a). Both CS-shFoxP2a and CS-shFoxP2i constructs resulted in equivalent downregulation of FoxP2 protein levels (p<0.0001 and p=0.0004, ANOVA), which were rescued in the presence of Cre recombinase (p=0.08 and p=0.93, ANOVA). (c) Validation of ICS constructs in vitro with western blotting. ICS constructs were generated and purified from in vitro cre recombination assay. Both CS and ICS constructs together with V5-tagged zebra finch FoxP2 were co-transfected into HEK cell as indicated. At 72 h after transfection, lysates of cells were subjected to immunoblotting with V5(to detect FoxP2) and tubulin antibody. In contrast to both CS-shFoxP2 constructs which are sufficient to downregulate FoxP2 protein level, neither ICS-shFoxP2 construct resulted in any change in the FoxP2 expression level. (d) Validation of CS constructs in the primary culture. CS-shScr or -shFoxP2 together with V5-tagged zebra finch FoxP2 were co-transfected into mouse cortical primary culture with or without Cre-GFP (+Cre & −Cre respectively). For both the CS-shScr and -shFoxP2 constructs, the expression of mCherry was maintained in the absence of Cre, whereas the expression of mCherry was turned off and BFP was turned on in the presence of Cre. Scale bar, 30 μm. (e) Representative parasagittal section shows the expression pattern of mCherry and tagBFP in Area X of an adult bird injected with CS-shFoxP2 and Cre-GFP constructs. Dashed lines outline the border of Area X. Scale bar, 100 μm.

**Figure S2.**
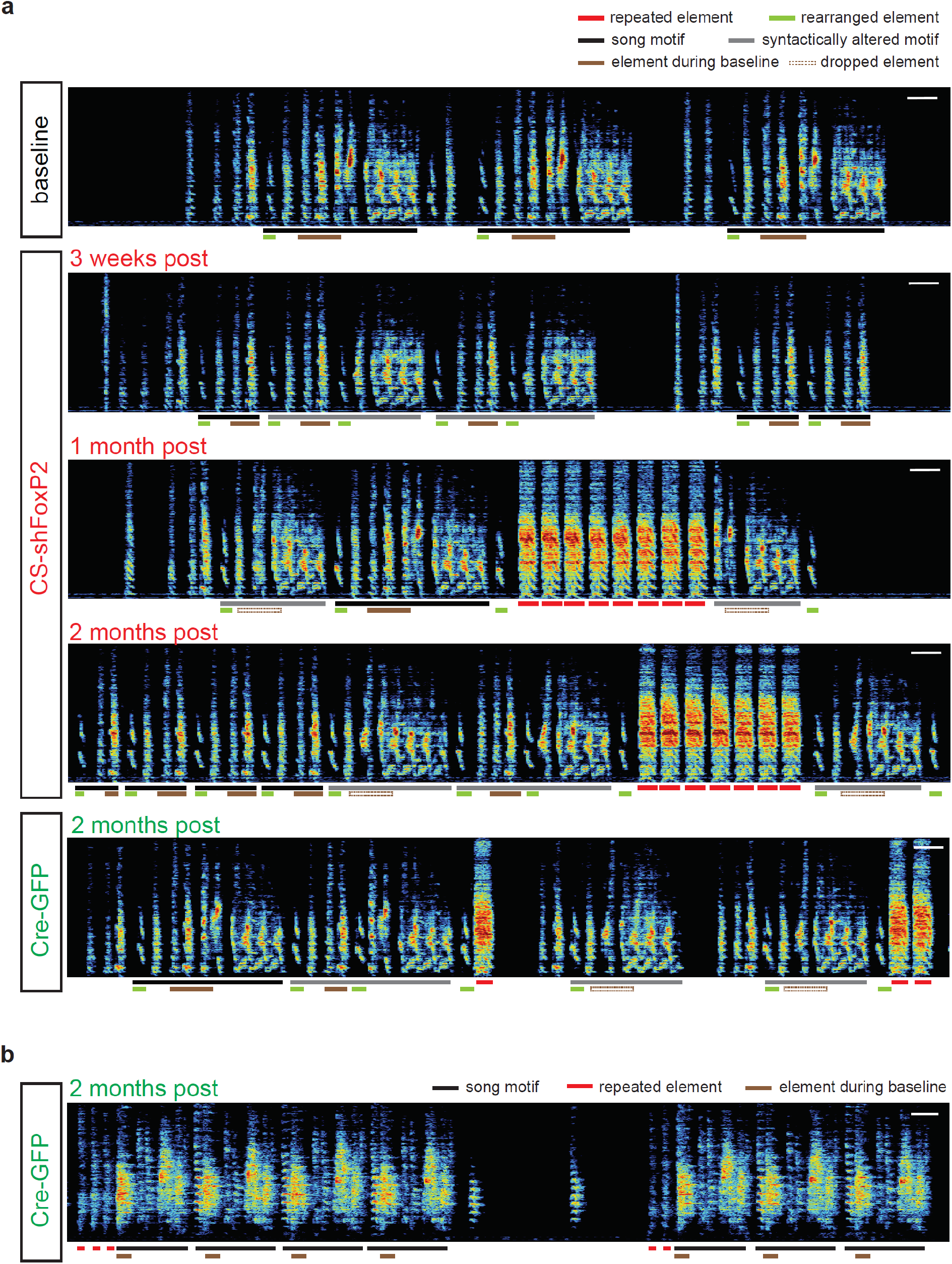
Changes in song following knockdown and/or rescue of FoxP2 in Area X. (a) Spectrograms of song recorded on the baseline day, 3 weeks, 1 month, and 2 months after injection of CS-shFoxP2 construct and 2 months after injection of Cre-GFP construct in Area X of an adult bird. A de novo syllable (red line) emerged in song bouts one-month post injection of the CS-shFoxP2 construct and the number of repetitions of this syllable was maintained for up to four months (data not shown) post injection of CS-shFoxP2 construct. In a subset of motifs, a portion of motif (brown and empty brown lines) was omitted one-month post injection of the CS-shFoxP2 construct. In another subset of motifs, the syllable immediately following that portion of the motif was replaced with the first syllable (green line) as early as three weeks post injection of the CS-shFoxP2 construct. The de novo syllable (red line) was retained in the end of a motif, whereas the number of repetitions in each song bout was significantly decreased 2 months following injection of Cre-GFP construct. All other changes in song caused by FoxP2 knockdown were not rescued 2 months following injection of Cre-GFP construct. Scale bar, 200 ms. (b) Spectrograms of one bout of song recorded two months after bilateral injection of Cre-GFP construct in Area X of adult birds which were previously injected with CS-shFoxP2 construct for 2 months (previous spectrograms were illustrated in Figure 2b). The number of repetitions of introductory elements in each song bout was restored to baseline level, and previous omitted syllable was fully recovered 2 months following injection of Cre-GFP construct

**Figure S3.**
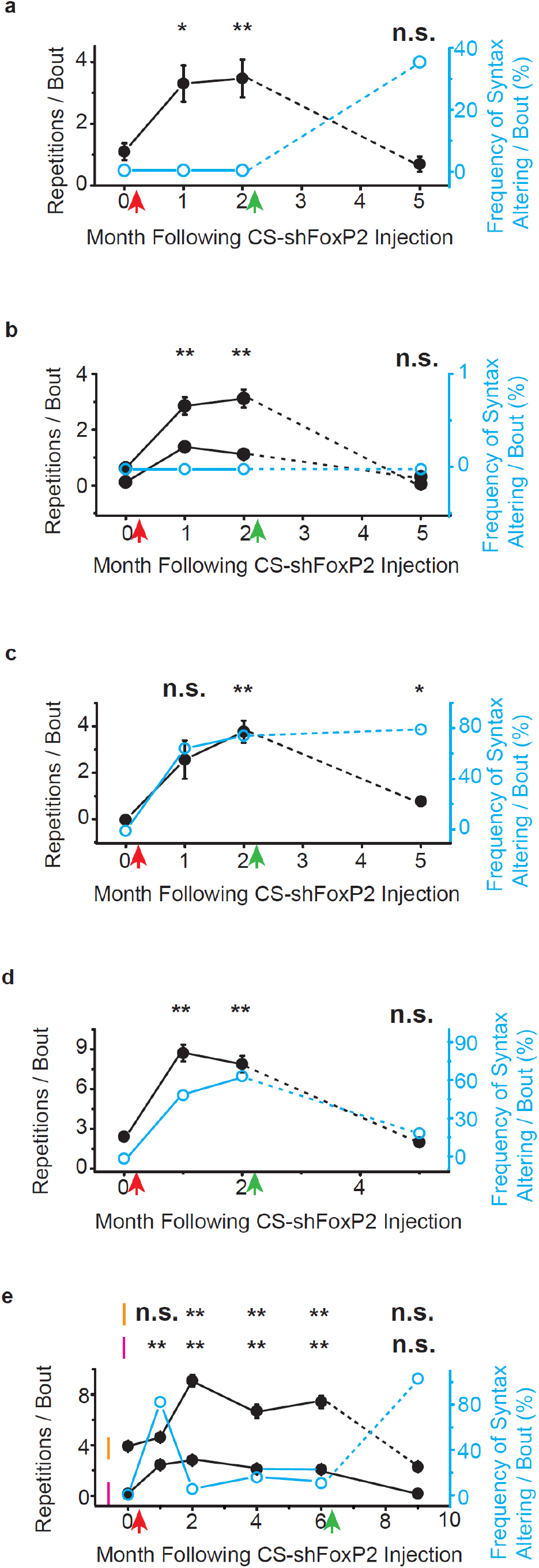
Reversal of FoxP2 knockdown in adult zebra finches rescues vocal repetitions but not changes in song syntax. Changes in the number of vocal repeats and frequency of syntax altering per song bout for five CS-shFoxP2+ birds (a-e, each panel represent an individual bird) who were injected with AAV encoding Cre-GFP (green arrow) 2-6 months after the initial knockdown of FoxP2(red arrow).

**Figure S4.**
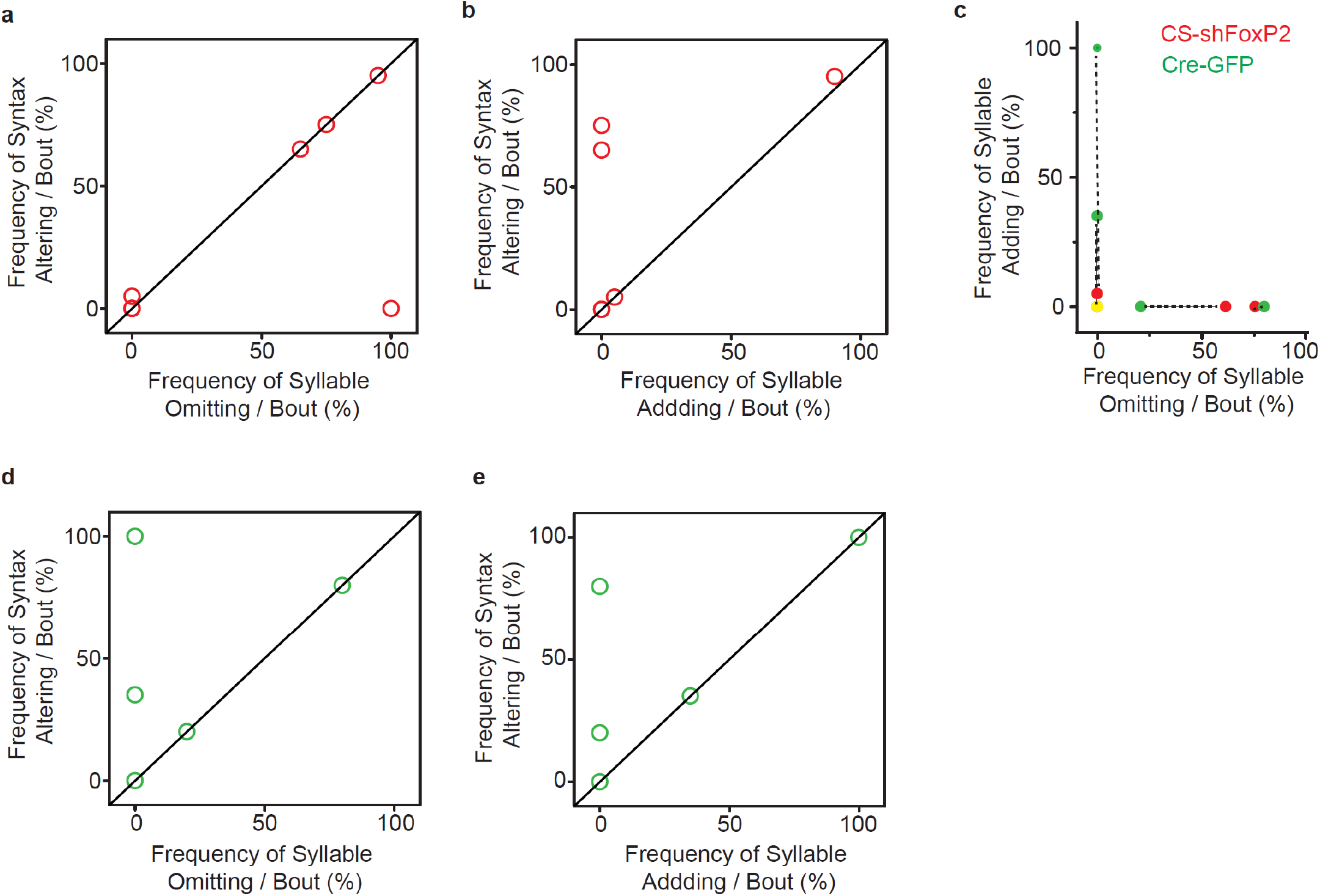
Relationship between syllable adding or omitting and syntax altering in CS-shFoxP2+ birds before or after reversal of FoxP2 knockdown. (a) The frequency of syllable omitting and syntax altering per song bout 2 months after injection of CS-shFoxP2 construct (n=8 birds, each circle represents an individual bird). (b) The frequency of syllable adding and syntax altering per song bout 2 months after injection of CS-shFoxP2 construct (n=8 birds, each circle represents an individual bird). (c) Changes in the frequency of syllable omitting and adding in CS-ShFoxP2+ birds(n=5) before (red circle, CS-shFoxP2+) and 2 months after injection of AAV encoding Cre-GFP (green circle, Cre-GFP+). Dashed line connects the same bird before and after reversal of FoxP2 knockdown. (a) The frequency of syllable omitting and syntax altering per song bout 2 months after injection of AAV encoding Cre-GFP in CS-shFoxP2+ birds (n=5, each circle represents an individual bird). (b) The frequency of syllable adding and syntax altering per song bout 2 months after injection of AAV encoding Cre-GFP (n=5, each circle represents an individual bird).

**Figure S5.**
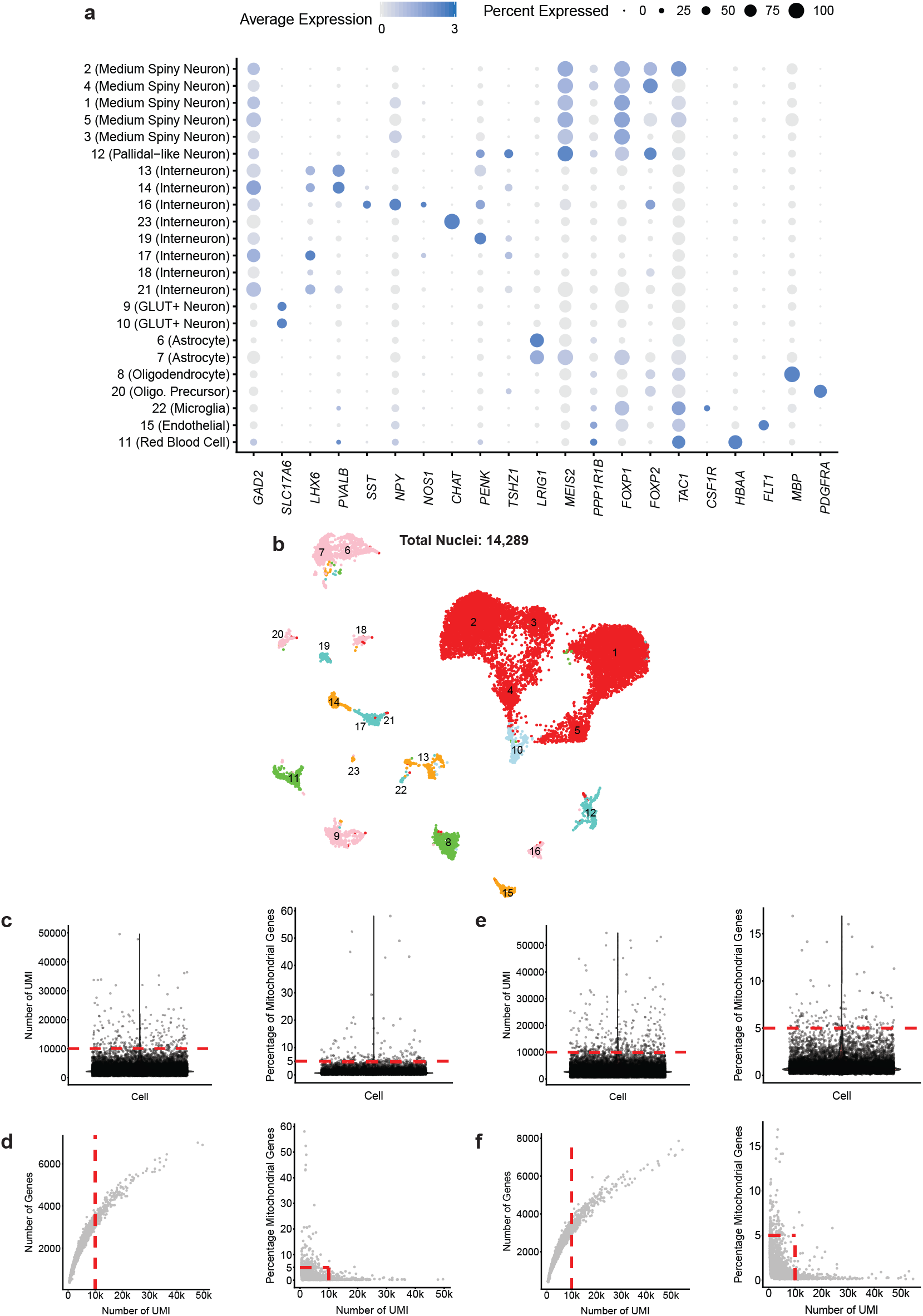
Details of single-cell transcriptomics. (a) A heatmap of normalized expression for genes used to assign identities to cell types for a combined analysis of the CS-shScr+ and CS-shFoxP2 datasets (corresponding to Figure 4a). Expression was normalized globally across all genes, but a different scale is shown for each gene based on the highest normalized value. (b) A UMAP projection of nuclei from Area X of an independent analysis of CS-shScr+ birds (not merged, corresponding to Figure 4c), and a plot of cell types by percentage of total nuclei. Clusters are numbered in ascending order by decreasing size (1-largest; 23-smallest). (c) For CS-shScr+, density plot of the number of UMIs per cell (left) and the percentage of mitochondrial genes in each cell (right). The analysis only included cells with UMI < 10,000 and < 5% mitochondrial genes (indicated by the red dashed line). (d) For CS-shScr+, a scatterplot of the number of UMI and the number of genes (left) or percentage of mitochondrial genes (right). Each dot is a cell. The cells within the red dashed box, corresponding to the filters in (c), were the cells analyzed. (e) For CS-shFoxP2+, density plot of the number of UMIs per cell (left) and the percentage of mitochondrial genes in each cell (right). The analysis only included cells with UMI < 10,000 and < 5% mitochondrial genes (indicated by the red dashed line). (f) For CS-shFoxP2+, a scatterplot of the number of UMI and the number of genes (left) or percentage of mitochondrial genes (right). Each dot is a cell. The cells within the red dashed box, corresponding to the filters in (e), were the cells analyzed.

**Figure S6.**
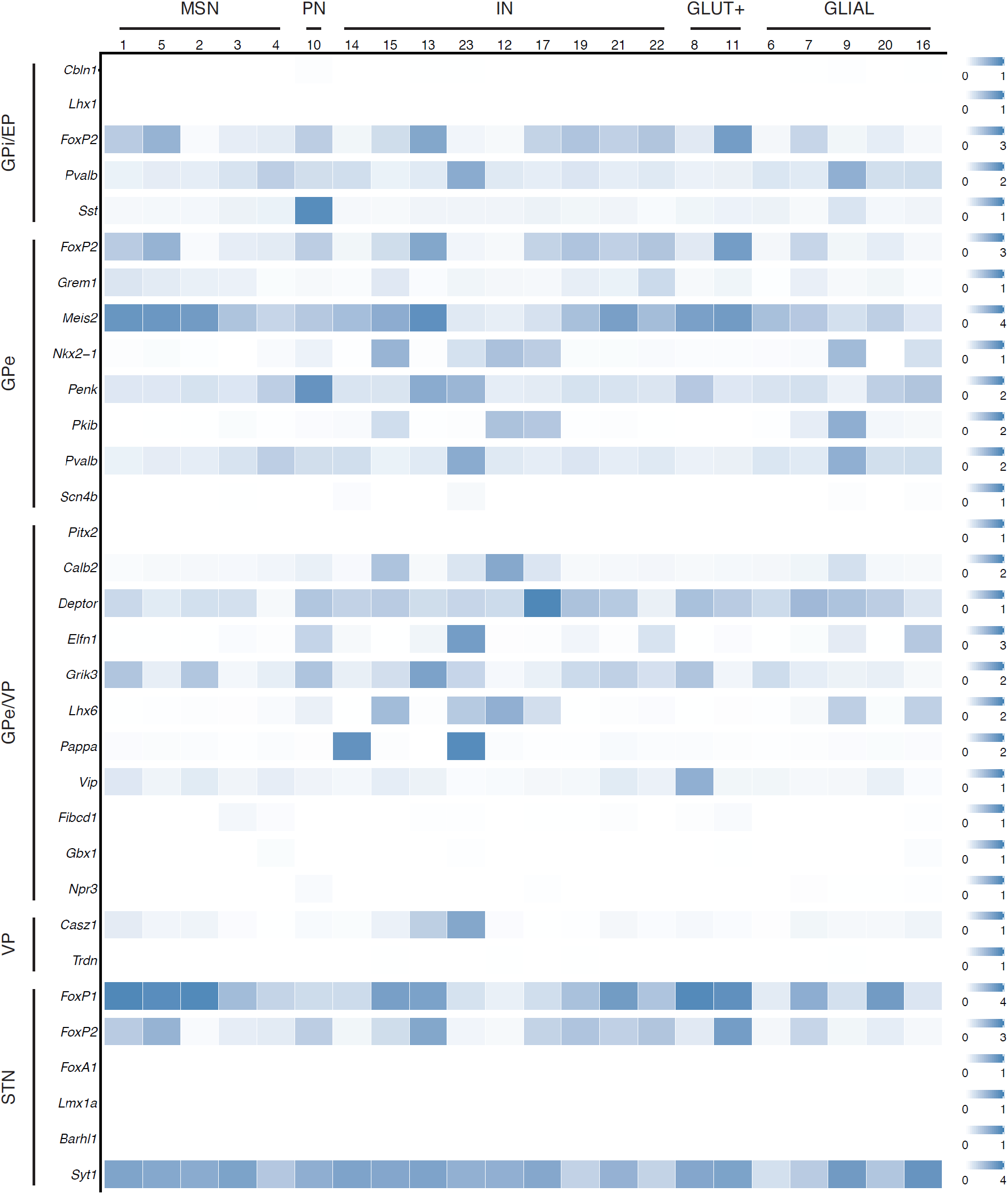
Supplemental gene markers of basal ganglia regions. As in Figure 4c, expression was normalized globally across all genes, but a different scale is shown for each gene based on the highest normalized value. Gene markers were selected from published studies (GPi/EP^88, 89^; GPe^88, 90^; GPe/VP^88^; VP^88^; STN^88, 91^).

**Figure S7.**
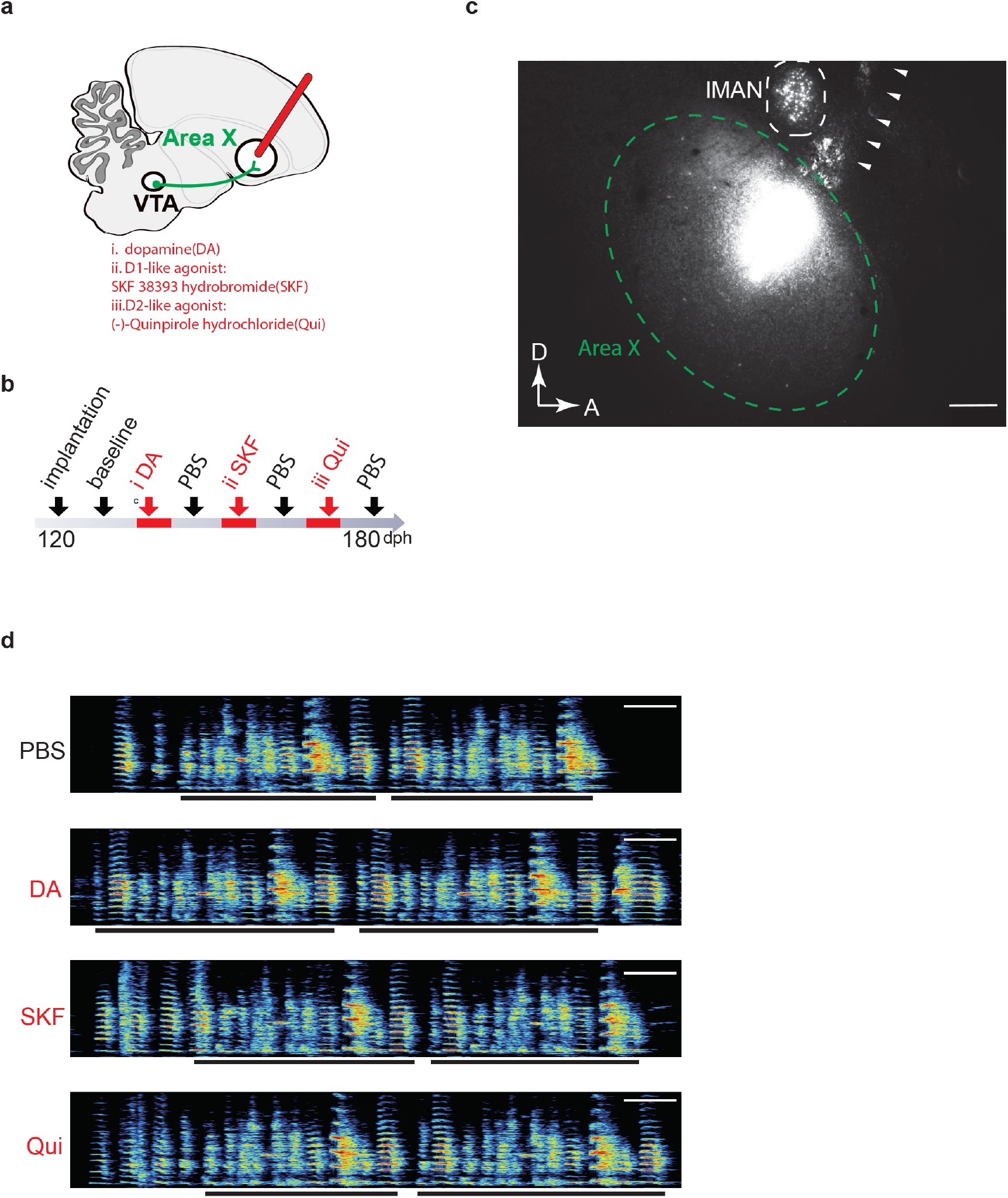
Lack of changes in vocal repetitions with pharmacological manipulation of dopamine or dopamine receptors. (a-b) Schematic of experimental design. Dopamine (DA, i), D1-like agonist: SKF 38393 hydrobromide (SKF, ii.) and D2-like agonist: (-)-Quinpirole hydrochloride (Qui, iii.) were infused bilaterally into Area X of behaving adult birds through a microdialysis system. Each chemical was delivered individually for 4-8 days, a week apart from each other. (c) Parasagittal section shows implantation track of microdialysis probe in Area X and anterior to lateral magnocellular nucleus of the anterior nidopallium (lMAN). A fluorescent retrograde tracer Fast blue was delivered to Area X by microdialysis probe in the end of each experiment to visualize field of pharmacological manipualtion. The fluorescent signals indicate the center of local infused Fast blue in Area X and retrograde labelled cells in lMAN, respectively. Triangles indicate the implantation track and dashed lines outline the border of Area X (in green) and lMAN (in white). Scale bar, 250 μm. (d) Spectrograms of song recorded from one adult bird implanted with microdialysis probes in Area X on the baseline day (infused with PBS) and third day during chronic infusion of DA, SKF and Qui. Black lines indicate motifs. Scale bar, 200 ms.

**Figure S8.**
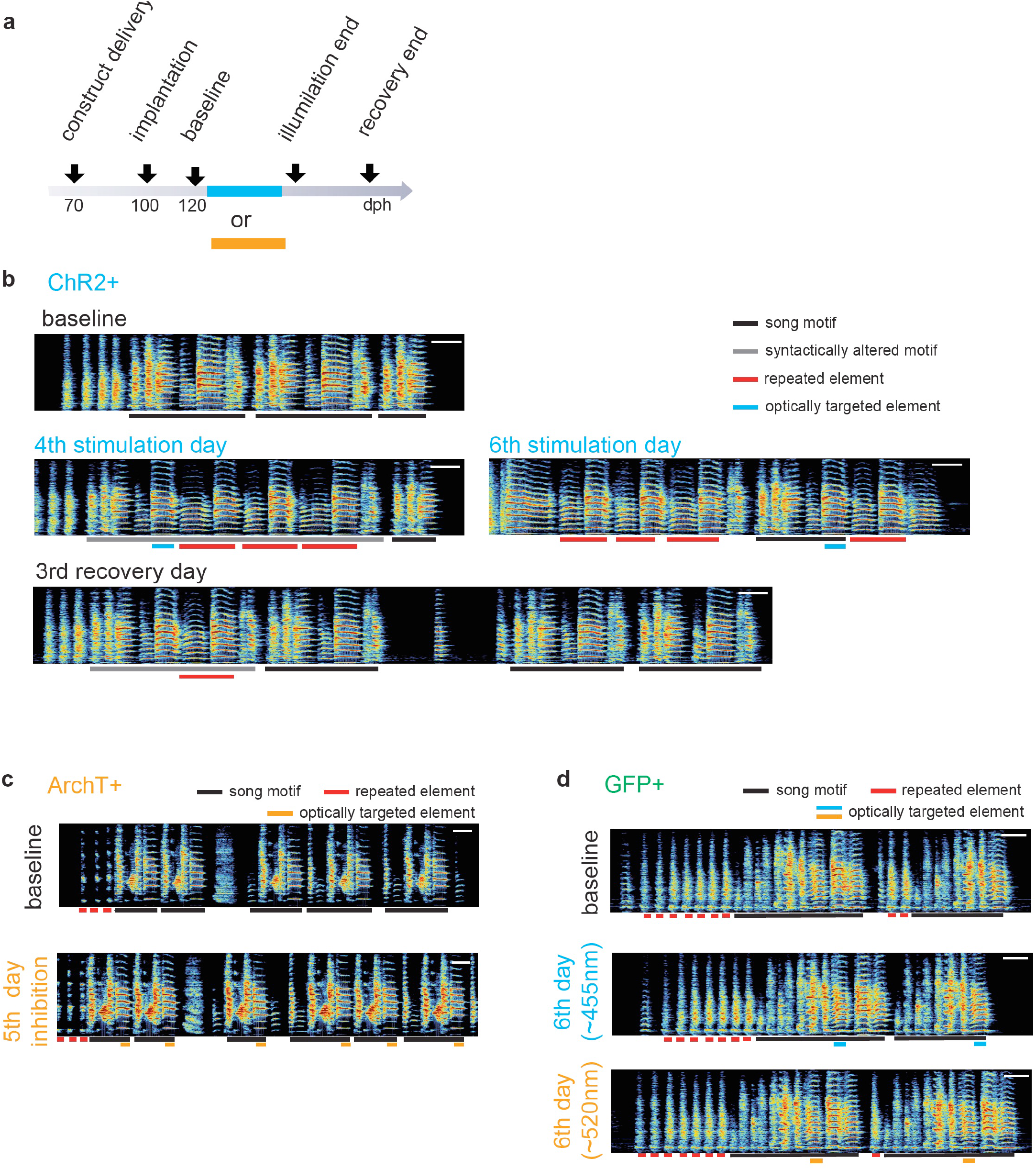
Song contingent optical manipulation of dopamine terminals in Area X. (a) Schematic of experimental design for optogenetic manipulation of dopamine release from VTA terminals. Optogenetic constructs were delivered into VTA in juvenile birds (~70 dph) and cannula pointing to Area X were not bilaterally implanted until the song crystallized(~100dph). Optogenetic manipulations began at least 1week post cannulas were implanted to allow birds to fully recover and to begin singing again (blue box, wavelength of LED ~455 nm; green box, wavelength of LED ~520 nm). Songs recorded before the illumination started and after the illumination ceased were referred as song of baseline and recovery, respectively. (b) Spectrograms of song bout recorded from a ChR2+ bird on the baseline day, 4th/6th stimulation day and 3rd recovery day illustrating a bird repeating a pair of syllables within the song motif or in the beginning or end of the song motif. Scale bar, 200 ms. (c-d) Lack of changes in vocal repetitions in birds expressing ArchT or GFP following optical manipulations. (c) Spectrograms of song recorded from a ArchT+ bird on the baseline day and 5th inhibition day. Light pulses (~520 nm, 100 ms) were delivered over the target syllable in a subset of variants during inhibition days. Scale bar, 200 ms. (d) Spectrograms of song recorded from a GFP+ bird on the baseline day(top) and 6th day of optical illumination. Light pulses (middle, blue light, ~455nm, 100 ms; bottom, green light, ~520 nm, 100 ms) were delivered over the target syllable (blue or orange line) in a subset of variants during illumination days. Scale bar, 200 ms.

